# Adaptive Functions of Structural Variants in Human Brain Development

**DOI:** 10.1101/2023.09.25.558917

**Authors:** Wanqiu Ding, Xiangshang Li, Jie Zhang, Mingjun Ji, Mengling Zhang, Xiaoming Zhong, Yong Cao, Xiaoge Liu, Chunqiong Li, Chunfu Xiao, Jiaxin Wang, Ting Li, Qing Yu, Fan Mo, Boya Zhang, Jianhuan Qi, Jie-Chun Yang, Juntian Qi, Lu Tian, Xinwei Xu, Qi Peng, Wei-Zhen Zhou, Zhijin Liu, Aisi Fu, Xiuqin Zhang, Jian-Jun Zhang, Yujie Sun, Baoyang Hu, Ni A. An, Li Zhang, Chuan-Yun Li

## Abstract

Quantifying the structural variants (SVs) in nonhuman primates could provide a niche to clarify the genetic backgrounds underlying human-specific traits, but such resource is largely lacking. Here, we report an accurate SV atlas in a population of 562 rhesus macaques, verified by two public SV benchmarks, an inhouse benchmark of eight macaque genomes with long-read sequencing and another inhouse benchmark of one macaque genome with whole-genome assembly. This accurate, quantitative SV map indicates stronger purifying selection on inversions, one type of poorly-clarified SVs to date, especially for those located on regulatory regions, suggesting a strategy for prioritizing inversions with the most important functions. Based on the distribution and the evolutionary features of these inversions in macaque population, we then identified 75 human-specific inversions, clarified their functional effects and prioritized them. Notably, the top-ranked inversions have substantially shaped the human transcriptome, through their dual-effects of reconfiguring the ancestral genomic architecture and introducing regional mutation hotspots at the inverted regions. As a proof-of-concept, we linked *APCDD1*, located on one of these inversions with the highest rank score and downregulated in human brains, to neuronal maturation. The accumulation of human-specific mutations on its promoter region, accelerated by the formation of the inversion, contributed to the decreased expression in humans. Notably, the overexpression of *APCDD1* could accelerate the neuronal maturation, while its depletion in mice delays the neuronal maturation. This study thus highlights the contribution of SVs, especially the inversions, to the distinct features in human brain development.

## Introduction

Genetic variation is a source of genetic novelty in shaping population structures and species-specific traits(1–8), which could be divided into categories based on their sizes, ranging from single-nucleotide variants (SNVs) to large-scale structural variants (SVs). SVs can be further classified into two categories: balanced SVs and unbalanced SVs(9). Unbalanced SVs are accompanied by gains or losses of DNA fragments, such as deletions and duplications, whereas balanced SVs can cause chromosomal rearrangements, such as inversions and translocations. Considering their larger sizes, SVs are expected to have stronger effects on transcription regulation than SNVs and short insertion/deletion (indel) variants(10). However, despite a growing awareness of their significance, the characterization and functional interrogation of SVs have largely lagged behind that of SNVs, partially due to the technical challenges in accurately identifying SVs with short reads obtained *via* the next-generation sequencing. Moreover, the public benchmarks for the evaluation of the performance of SV detection are limited to a small number of validated SVs in human samples(11–13), further hindering the studies of complex SVs and those in other species.

Rhesus macaque (*Macaca mulatta*) is a non-human primate species closely related to humans in terms of the genome sequences and the physiology(14, 15). Quantification of SVs in macaque populations could thus promote the understanding of their features, turnover and evolutionary significances, and further provide a niche to clarify the genetic backgrounds underlying human-specific traits. Recent advances in the assembly of macaque reference genomes and the generation of a batch of genome resequencing data have facilitated the profiling of genetic variants in macaque populations(6, 15–20). However, the data are scattered throughout the literatures and are largely generated with short-read sequencing, which is error-prone to be used in SV calling with standardized algorithms. In addition, the deep integration of these data from multiple subpopulations of macaques and the issue of kinships among these animals are not well addressed, further confounding the in-depth population genetics studies. Overall, the challenges in the deep integration of these confounded and scattered macaque population genomics data, the difficulty of accurate identification of SVs with short-read sequencing, and the lack of comprehensive benchmarks for macaque SV evaluation have hindered the clarification of the functions and evolutionary significance of SVs in primates.

Here, we provide the largest and accurate macaque SV atlas to date by integrating WGS data from 1,026 macaques. We addressed the issue of kinships using genome-wide SNV profile, developed an accurate pipeline for SV calling, and established a three-tier benchmark for comprehensive SV evaluation. Furthermore, we explored the evolutionary turnover of these SV events and proposed a practical strategy for prioritizing those with the most important functions in shaping human adaptive evolution. Based on this, we identified 75 human-specific inversions and prioritized those inversions which have substantially shaped the human brain transcriptome.

## Results

### Definition of a well-annotated population of independent macaques

To define a clean population of independent macaques for unbiased population genetics analyses of SVs, we first attempted to establish a comprehensive SNV atlas of macaques, which could provide a genetic basis to examine their identity information, such as subpopulations, sexes and kinship relationships. To this end, we first constructed a better reference macaque genome through the integration of three recent genome assemblies based on the third-generation sequencing. As Mmul_10 (rheMac10) represents the genome assembly with the highest integrity in sequence continuity and base accuracy (**Figs. S1A** and **B**), we used this assembly as the template and filled gaps by aligning with it the sequences of two other genome assemblies from an Indian-origin macaque (rheMac8) and a Chinese-origin macaque (rheMacS). A total of 40 gaps in Mmul_10 were filled with the corresponding sequences from the two genomes and further evaluated with the high-coverage Bionano optical map from a macaque and PacBio long-read sequencing from eight additional macaques (**Figs. 1A**, **S1 and S2**, **Table S1–S3**, **Materials and Methods**). The results indicate that these gaps should represent real genomic gaps in Mmul_10 assembly, rather than the assembly errors from other macaque genome assemblies, or individual-specific structural variants. Notably, the 40 gap regions that were closed here were distributed across the genome spanning 721,984 bp, and some were located in regions with high gene density (**Fig. 1A**). We named the genome with filled gaps “rheMac10Plus” and performed subsequent analyses with this improved reference genome.

**Fig. 1.**
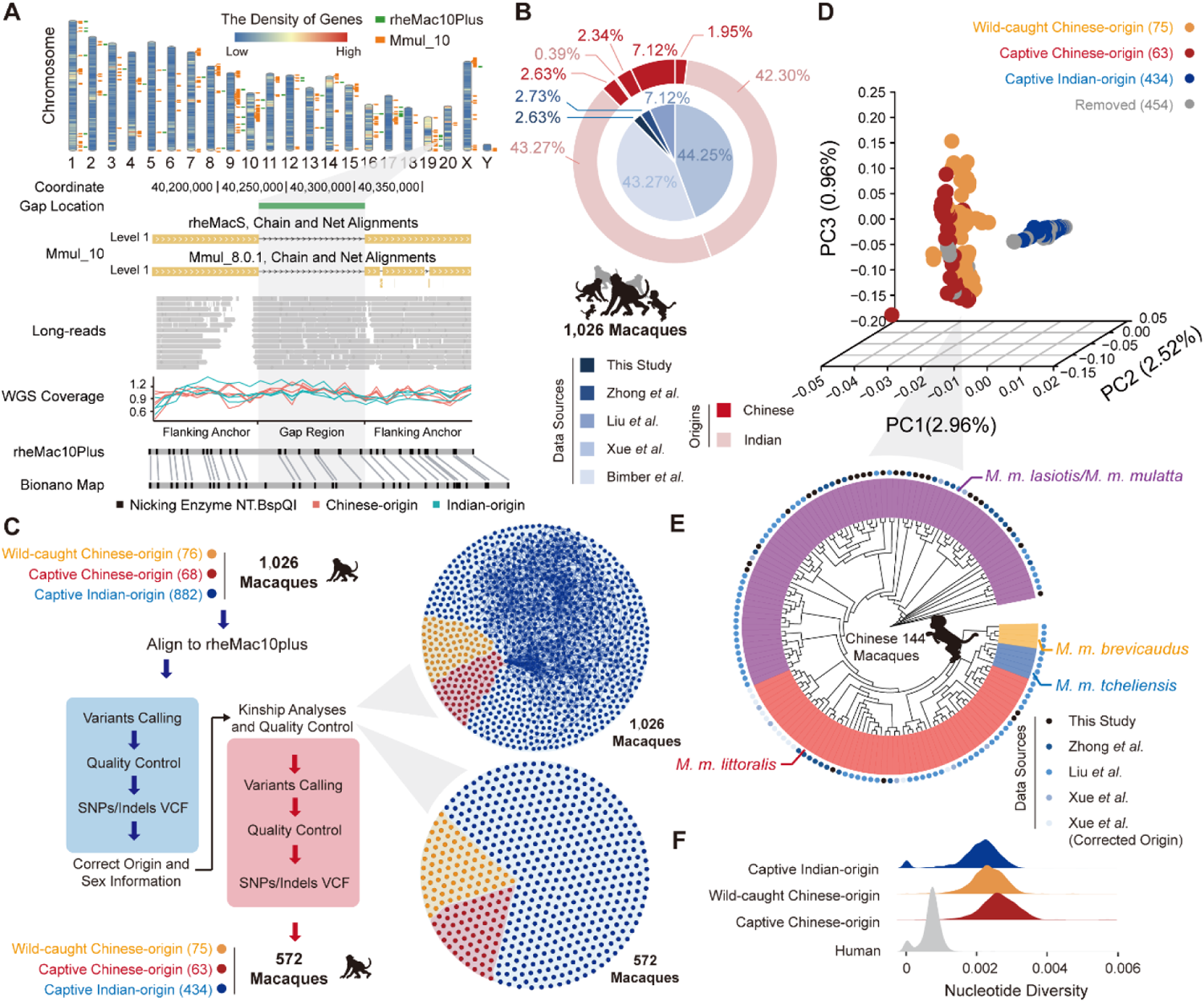
Population genetic landscape of 1,026 macaque genomes. (**A**) Chromosome karyotype showing 40 filled gaps, as indicated by green bars in the rheMac10Plus assembly. The density of genes across the genome is shown in the heatmap. For one of the filled gaps on chromosome 19, the Bionano optical map of one macaque, the long reads of eight macaques, and the coverage of the short reads of ten macaques (yellow: Chinese-origin macaques; green: Indian-origin macaques) were aligned and shown accordingly. (**B**) Sources (inner layer) and geographic origins (outer layer) of the 1,026 macaques. (**C**) Schematic diagram of the workflow for variant calling with a two-round strategy (blue: first round of calling; red: second round of calling). The original set of macaques (1,026 animals) and the set after quality control (572 animals) were partitioned into three clusters (red: captive Chinese-origin macaques; yellow: wild-caught Chinese-origin macaques; blue: captive Indian-origin macaques) based on their genetic profiles. The pairs of macaques with significant kinship relationships are linked by lines. (**D**) Three-dimensional PCA plot showing the relationships of the 572 macaques according to SNV genotypes. (**E**) Neighbor-joining tree showing the genetic distance of the Chinese-origin macaques. Different data sources are indicated by colored dots in the outer layer. Yellow: *M. m. brevicaudatus*; blue: *M. m. tcheliensis*; orange: *M. m. littoralis*; purple: *M. m. lasiotis* or *M. m. mulatta*. (**F**) The genome-wide distribution of nucleotide diversity of captive Indian-origin macaques (blue), captive Chinese-origin macaques (red), wild-caught Chinese-origin macaques (yellow) and humans (gray).

Based on the improved macaque genome, we performed whole genome sequencing of 27 captive Chinese-origin macaques and achieved an average coverage of 32-fold (**Table S4**). We also integrated public genome resequencing data of 41 captive Chinese-origin macaques, 76 wild-caught Chinese-origin macaques, and 882 captive Indian-origin macaques (**Fig. 1B**, **Table S5**)(16–19). Overall, the genome resequencing data of a total of 1,026 macaques were obtained (**Fig. 1B**), in which 862 out of 1,026 (84.0%) were sequenced at a depth of more than 30-fold (**Table S5**). The resequencing data of the 1,026 macaques were then aligned to the rheMac10Plus genome and further subjected to a two-round variant calling (**Fig. 1C**, **Materials and Methods**). A total of 81.3 million SNVs were then identified.

On the basis of this genetic profile, we then carefully examined the identity information of these macaques to remove animals that would confound the conclusions of the subsequent population genetic analyses of SVs (**Table S5**). Notably, through a principal component analysis (PCA) of the SNV genotypes across the 1,026 macaques, we found that 12 animals initially identified as having an Indian origin were actually of Chinese origin (**Fig. S3A**). We then examined the reported sex information of these animals by checking the read density along the X chromosome, and found that four female macaques were falsely recorded as male, while one male macaque was reported as a female animal (**Fig. S3B**). We further assigned the information for 74 wild-caught Chinese-origin macaques with missing sex information (**Fig. S3C)**. Finally, we carefully examined the kinship of these macaques according to their genetic profiles and subsequently removed 454 macaques that were closely related to other macaques (**Fig. 1C**, **Materials and Methods**). Overall, after stringent filtering steps, 572 independent macaques were retained in the following analyses, including 434 captive Indian-origin macaques, 63 captive Chinese-origin macaques and 75 wild-caught Chinese-origin macaques (**Table S5**).

The PCA analysis of the 572 independent macaques revealed distinct divergence between the Indian-origin and Chinese-origin macaques (**Fig. 1D**), while the captive Chinese-origin macaques were indistinguishable from the wild-caught Chinese-origin macaques (**Fig. 1D**), indicating a weaker effect of domestication on the genetic backgrounds of macaques in comparison to the effect of geography. As the wild-caught Chinese-origin macaques with definite habitat information could be divided into five subpopulations widely used in biomedical studies(18), we assigned subpopulation information to other Chinese-origin macaques by constructing a neighbor-joining phylogeny combining the two groups of macaques, in which 72 additional Chinese-origin macaques were annotated with the subpopulation information accordingly (**Fig. 1E**, **Materials and Methods**).

Overall, we defined a well-annotated population of 572 independent macaques, resulting in a macaque SNV profile with a total of 79.6 million SNVs and 9.1 million indels. Consistent with previous findings(16, 19), we observed a comparable transition-to-transversion ratio, and significantly increased nucleotide diversity in both Indian-and Chinese-origin macaque populations in comparison to that in humans (**Fig. 1F**). This population thus represent a clean population for unbiased population genetics analyses of SVs.

### Construction of the macaque SV map

Based on the resequencing data of 572 independent, well-annotated macaques, we next attempted to identify SV events at the population scale. Notably, it is error-prone to use standardized tools to identify SVs with short reads. To identify an accurate SV map, we then developed a new pipeline by carefully adjusting the parameters of each tool, performing meta-analyses to integrate the results of different tools, and introducing stringent filters to control for false-positives (**Fig. 2A**, **Materials and Methods**). In such a case, the requirements to define an SV event are more stringent than previous practices. As a note, the parameters were set and adjusted according to a benchmark of two macaques with both short-read resequencing data and the genome assemblies available (Mmul_10 and rheMacS). Especially, in this pipeline, the signatures of paired-end reads and split reads were adequately integrated to capture candidate regions of inversions (**Fig. 2A**), a type of balanced SVs that is difficult to detect using short reads. Although most of the reads are not informative in indicating the position and boundaries of them, the split reads and the poorly-paired reads in paired-ends sequencing are informative to indicate the inversion event. It is thus practical to pinpoint these events with high-coverage short reads, as long as the two types of informative reads are adequately analyzed. As a proof-of-concept, we included an example for one inversion we identified to illustrate the principle in calling inversions with both short reads and long reads (**Fig. 2B**).

**Fig. 2.**
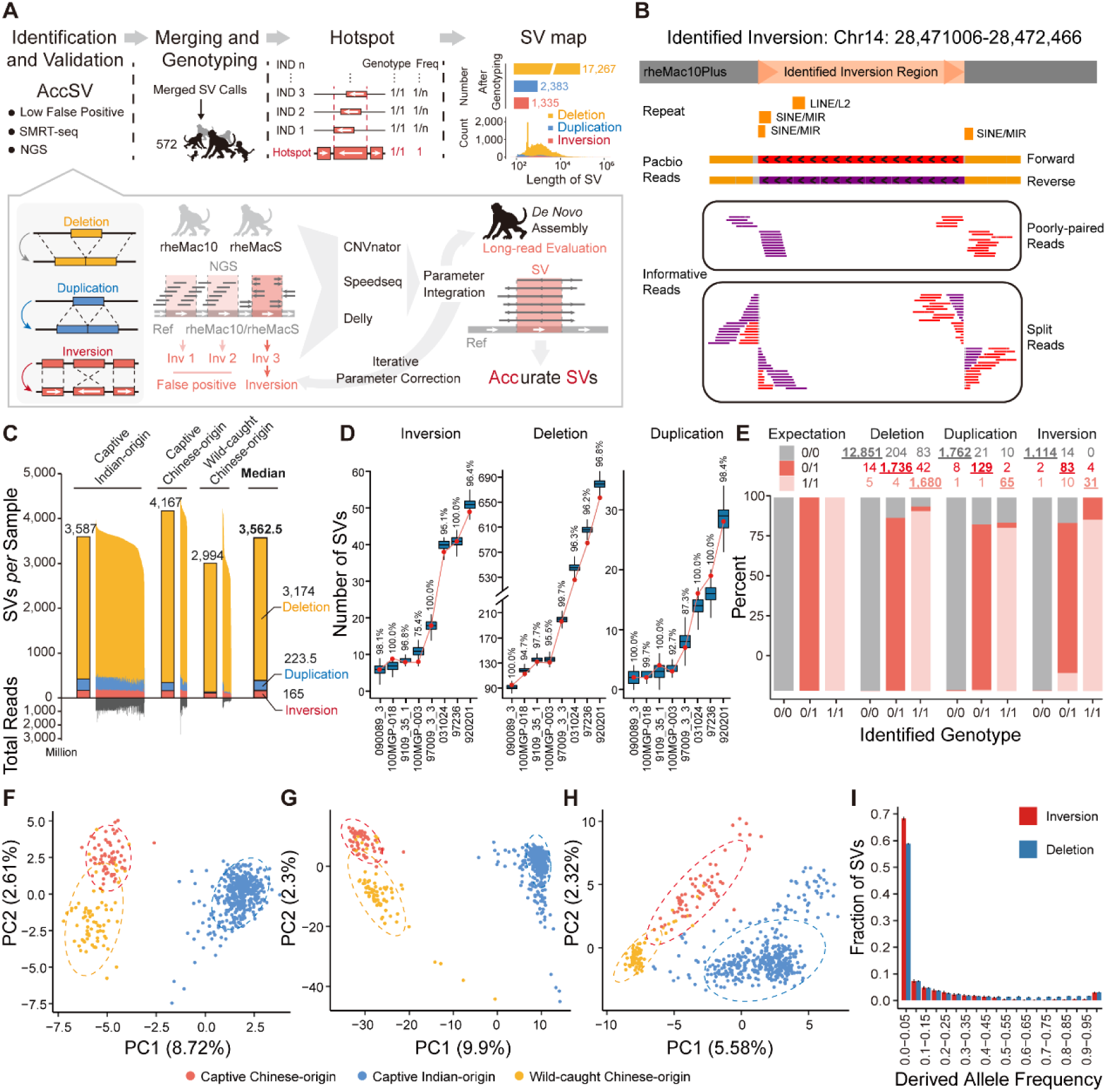
Construction and characterization of SV map for macaque population. (**A**) The pipeline for SV map construction, including the processes of SV identification, validation, genotyping, SV hotspot definition and allele frequency calculation. (**B**) The genomic region of one inversion we identified was shown as an example, with the split reads, poorly-paired reads and the long reads supporting its existence aligned and shown accordingly. (**C**) The distribution of the count of SVs *per* macaque genome for three types of SVs in macaques of different origins. The median number of SVs is shown for each group. The total number of reads of deep sequencing for each macaque is also shown. (**D**) Verification of the SV events with long HiFi reads of eight macaques. Boxplots showing the distribution of the theoretical number of verified SVs at the current sequencing depth of HiFi reads, obtained from 10,000 times of simulations. The detected number of verified SVs in each macaque was indicated by the red dot. (**E**) Validation of the genotypes of SVs in one macaque based on the long-read sequencing and genome assembly in one macaque. For each type of genotype identified with short reads (0/0, 0/1, or 1/1), the percentages of verified SVs are summarized and shown in different colors. The numbers of SVs of each type are shown, and those with verified genotypes are underlined. (**F-H**) PCA plots showing the relationships of the 562 macaques according to the genotypes of inversion (**F**), deletion (**G**) and duplication (**H**) variations. Macaques with different origins are labeled with different colors. (**I**) Site frequency spectra of the derived alleles for inversions and deletions in 562 macaques.

After identifying the SVs of each macaque, we then merged the SV calls and genotyped each SV in all of the 572 macaques (**Fig. 2A**). Briefly, considering the difficulties in defining SV breakpoints in each macaque at a single-nucleotide resolution, which may lead to the underestimation of the SV allele frequency, we defined “SV hotspot” indicating the consensus SV regions shared by different macaques with slightly different boundaries (**Fig. 2A**, **Materials and Methods**). Large SVs with lengths over 10 Mb were removed to avoid false positives in SV identification with short reads, and ten macaques with abnormally high numbers of SV hotspots were eliminated from the subsequent analyses (**Materials and Methods**). Finally, a total of 20,985 SV hotspots were defined for the remaining 562 macaques, including 1,335 inversions, 17,267 deletions and 2,383 duplications. As a note, the distance between two adjacent SV hotspots across the genome (median distance: 1,284,360 bp for inversions; 96,398 bp for deletions; 743,330 bp for duplications) are typically much larger than the size of these SV hotspots (median size: 1,025 bp for inversions; 954 bp for deletions; 330 bp for duplications), indicating that these hotspots could be clearly distinguished from neighboring hotspots. Using this atlas of SV hotspots as a reference, we further genotyped the SVs in each macaque and calculated the allele frequency for each SV hotspot in the population, which substantially increased the sensitivity of SV calling in comparison to the previous round of *de novo* calling. Overall, a median of 3,562 SVs were detected in each macaque, including 165 inversions, 3,174 deletions and 223 duplications (**Fig. 2C**).

To confirm that these events represented *bona fide* SVs, we then developed a three-tier benchmark to evaluate the performance of our pipeline in SV calling. We first evaluated the pipeline with public benchmarks in human. Briefly, we selected the HG002 genome with verified deletion calls as the benchmark callset(21), and evaluated the performance of our pipeline in calling these deletion events. Notably, the pipeline achieved a high precision score (97.8%). We then used another benchmark callset of HG001 genome from Pendleton *et. Al.*(22) to evaluate the performance of our pipeline in calling inversions. The pipeline achieved a precision score of 88.9% in calling these inversions. As the parameters were adjusted according to the features of macaque short-read sequencing data, the real precision score in calling SVs in macaque populations should be higher.

Considering the complexity of SVs and the lack of comprehensive, high-quality reference SV standards in macaques, we further developed in-house benchmarks for evaluating the SV calling in macaques, according to the principles of public benchmarking standards, in that the long sequencing reads, especially the *de novo* genome assembly, could provide a more accurate SV atlas. To this end, from the population of 562 macaques, we selected eight macaques and sequenced their genomes with different coverages of long HiFi reads (**Table S3**, **Materials and Methods**). For each macaque animal, we then evaluated the performance of our pipeline in calling deletions, inversions and duplications, using SVs called in this macaque by long HiFi reads as a benchmark. As the sequencing of these samples was not saturated, for each SV type in each macaque animal, we performed a simulation strategy to estimate the theoretical number of verified SVs at the current sequencing depth, assuming that the SVs and their genotypes were accurately identified with short reads (**Materials and Methods**). Overall, the average verification rates (97.1%, 95.2% and 97.3% for deletions, inversions and duplications, respectively) indicate the high accuracy of our pipeline in SV calling with short reads (**Fig. 2D**).

Finally, for another Chinese-origin, male rhesus macaque with normal phenotype from the population of 562 macaques, we sequenced its genome with high coverage long-read sequencing, and then *de novo* assembled its genome, which was further used to evaluate the performance of our pipeline in SV calling with short reads (**Materials and Methods**). Specifically, we applied SMRT long-read sequencing technology and sequenced the genomic DNA. 290 Gb of data were generated, with the N50 of the subread length of 14.3 kb and an average genome coverage of 96.9-fold (**Table S6**, **Materials and Methods**). We further *de novo* assembled its genome on the basis of the integration of the short-read sequencing, long-read sequencing and Bionano optical data, resulting a genome assembly with 2.99 Gbp informative bases, supported by 4,742 contigs (N50 = 4.6 Mbp, **Materials and Methods**). Based on the long reads and the assembled genome, we then evaluated the genotype of each SV identified with short reads of the same macaque (**Fig. 2E**). The verification rate (92.3%, 90.1% and 86.4% for deletions, inversions and duplications, respectively) indicates that the SVs we identified with short reads are accurate. For each type of SV events, one verified case was shown in **Fig. S4**.

In this accurate SV atlas of 562 macaques, we found similar densities of SVs across macaques with different origins and sexes, a pattern consistent with that in human subpopulations(3) (**Figs. 2C** and **S5A**). Furthermore, although the number of SVs located on each chromosome was correlated with the length of the chromosome, a pattern consistent with previous reports (**Figs. S5B–D**)(3, 23), we observed enrichments of inversions on chromosomes 16 and 19, and a depletion on chromosome 18; an enrichment of deletions on chromosome 5, and depletions on chromosomes 7 and X. These regions with unbalanced distributions of SVs need further investigations. Similar to the SNV profile (**Fig. 1D**), the profile of these SVs could discriminate macaques from different subpopulations with an even higher discrimination efficiency (**Figs. 2F–H**). Of a note, the genetic distances between the captive Chinese-origin macaques and the wild-caught Chinese-origin macaques were smaller than that between the captive Chinese-origin macaques and the captive Indian-origin macaques, recapitulating the above conclusion based on SNVs for a weaker effect of domestication on the genetic background in comparison to that of geography (**Figs. 1D**, **2F–H**).

### Genomic inversions are selectively constrained in macaque populations

To investigate whether SVs are selectively constrained, we inferred the ancestral state of each SV and compared the site frequency spectra of the derived allele for each type of SV events (**Materials and Methods**). Previous studies have reported that deletions are more deleterious than duplications(24). Notably, we found that the allele frequency spectrum of inversions was more left-skewed than that of deletions, indicating even stronger purifying selection on the fixation of inversions than deletions (**Fig. 2I**). This finding is consistent with previous reports that polymorphic inversions are largely deleterious due to recombination suppression and the subsequent accumulation of deleterious mutations(25–28). Considering the possibly stronger effects of inversion, we next focused specifically on this type of SV in our subsequent evolutionary and functional genomics analyses (**Discussion**).

We first investigated whether inversions with different features or genomic locations were shaped by natural selection to different degrees (**Fig. 3A**). When inspecting the distribution of these inversions, we found that they tended to be depleted in functional regions, such as exons (Permutation test, P value < 0.001), putative promoters (P value < 0.001) and enhancers (P value = 0.02; **Fig. 3B**), as significantly fewer events were located in these regions than in randomly-shuffled, length-matched regions used as a background. In contrast, these inversions were overrepresented in intergenic regions (P value < 0.001; **Fig. 3B**). Accordingly, the inversions in exonic regions showed an excess number of low-frequency variants of the derived allele, in comparison to those located on intronic regions (Wilcoxon rank-sum test, P value < 2.2e-16) or intergenic regions (P value < 2.2e-16; **Fig. 3C**).

**Fig. 3.**
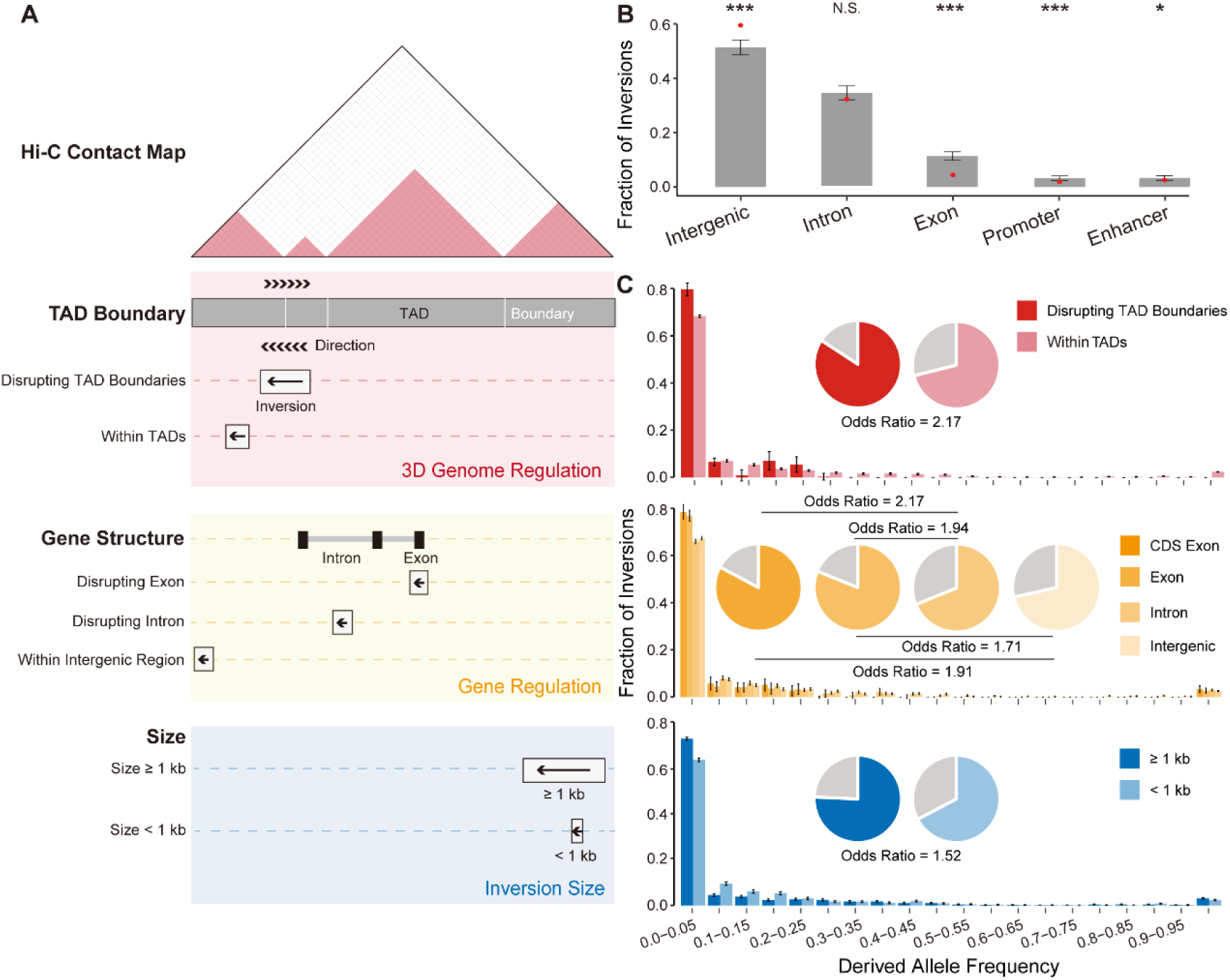
Inversions in regulatory regions are selectively constrained. (**A**) Classification of inversions by different features and genomic locations, including the sizes of the inversions, their locations on the genes and their three-dimensional genomic architecture. (**B**) Proportions of inversions at different genomic locations. The background distribution of inversions located in each genomic region, as estimated based on 1,000 shuffled regions with matched lengths, is shown in a bar plot, with the error bars representing the standard deviations. For each bar plot, the observed value is indicated as a red dot, with the empirical P value calculated as the percentage of the 1,000 replicates. *P value < 0.05, ***P value < 0.001, N.S., not significant. (**C**) Site frequency spectra of the derived allele for different classifications of inversions. For each group of inversions, the fraction of inversions with a low frequency of derived alleles (less than 5%) is shown and compared, with the odds ratios shown accordingly.

When further dividing these inversions into groups according to their sizes and locations on the three-dimensional genome (**Fig. 3A**), we found that the inversions with larger sizes showed an excess proportion of low-frequency variants compared with short inversions (Wilcoxon rank-sum test, P value < 2.2e-16; **Fig. 3C**). Moreover, inversions disrupting TAD boundaries, as defined in macaque fetal brains(29), also showed an excess of low-frequency variants, in that the frequency spectrum of the derived allele was significantly left-skewed relative to that of the inversions located within TADs (Wilcoxon rank-sum test, P value < 2.2e-16; **Figs. 3C** and **S6**).

We then investigated the distribution of the inversions with extremely low frequencies in various genomic regions. In contrast to inversions with higher frequencies, low-frequency inversions were typically larger in size (odds ratio = 1.52), and located with a higher proportion on gene regions (odds ratio = 2.17) or regulatory regions disrupting TAD boundaries (odds ratio = 2.17; **Fig. 3C**). These findings thus indicate that the inversions located in regulatory regions are subjected to stronger purifying selection, suggesting a practical strategy for prioritizing them with the most important functions, as the fixed inversions with stronger effects on gene structure or expression regulation were more likely maintained by selective pressure owing to their adaptive functions.

### Identification of 75 fixed human-specific inversions

Prompted by the assumption that the list of fixed inversions with stronger effects on gene structure or expression regulation should be enriched with functional inversions driving species-specific traits, we next identified human-specific inversions based on the above macaque SV atlas and comparative genomics analyses in multiple outgroups. To this end, we first identified 1,972 species-specific inversions between humans and macaques, ranging in size from 51 bp to 69 Mb, through genome-wide alignment followed by intensive manual curation in the UCSC genome browser (**Fig. 4A**, **Table S7**, **Materials and Methods**). The number of species-specific inversions on each chromosome was correlated well with the length of the chromosome, consistent with the observation of the polymorphic inversions in macaque population (**Fig. S7**).

**Fig. 4.**
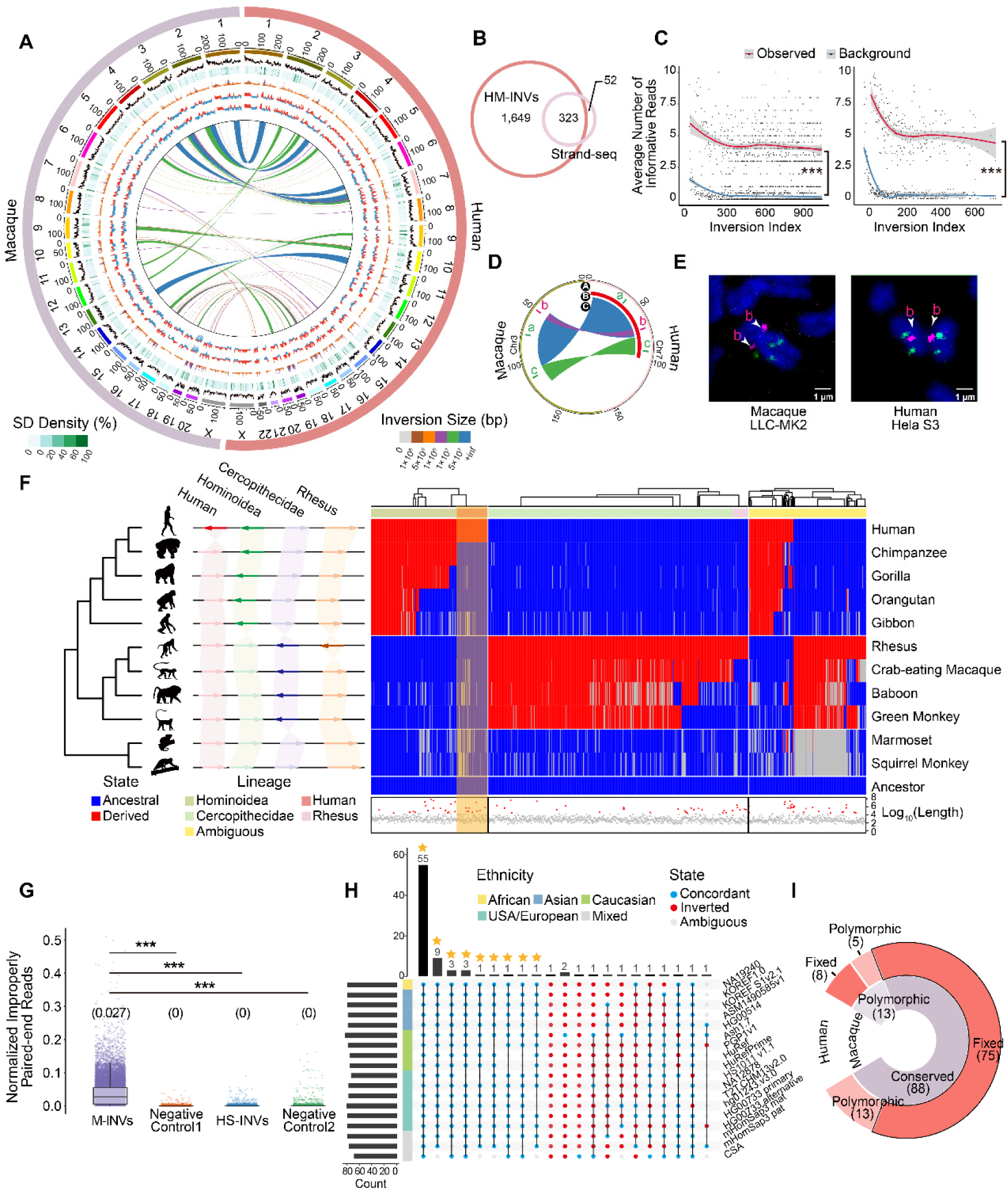
Identification of human-specific inversions. (**A**) Circos plot showing the profile of inversions across humans and macaques (HM-INVs) and the genomic features aligned according to the coordinates. From the outside to the inside: GC content (%), Segmental duplication (SD) density (%), gene density, A/B compartments from fetal cortical plates (CP) and germinal zone (GZ) and the locations of HM-INVs. The average GC contents are indicated by orange lines. Genomic regions with more than 10 genes are shown as purple bars in the gene density track. Tracks are plotted in 500 kb windows. (**B**) Overlap between HM-INVs in this study and the public list of inversions between humans and macaques as defined by in Maggiolini *et. al* with the Strand-seq(4) assay. (**C**) Validation of species-specific inversions between human and macaque with Strand-seq data. For candidate inversions with reads coverage ≥ 3 in the Strand-seq study, the average numbers of Strand-seq informative reads (**Observed**) were shown and compared with the background (**Background**, see details in **Materials and Methods**), for candidates identified specifically in our study (**left**), or candidates identified by both studies (**right**). Inversions were arranged in descending order of their length. Local regression curves for the average numbers of the informative reads (red) and the background (blue) were shown. Wilcoxon rank-sum tests, ***P value < 0.001. (**D**) Circos plot depicting the arrangement of one complex HM-INV chosen for FISH validation. The track A represents the genomic regions where the probes were designed, with the order of colors indicating the expected form of inversions in humans and macaques based on the definition in this study. Tracks B and C display the forms of these inversions identified by Strand-seq (one large inversion) and in this study (three complex inversions with breakpoint reuse), respectively. (**E**) Validation of the complex HM-INV in **D** in the macaque LLC-MK2 cell line (left) and human HeLa S3 cell line (right). (**F**) Schematic illustration (left) of four classes of lineage-specific inversions, including human-specific (**Human**), Hominoidea-specific (**Hominoidea**), Cercopithecidae-specific (**Cercopithecidae**) and macaque-specific (**Rhesus**) inversions. Heatmap (right) showing the arrangement of HM-INVs in comparison to the estimated ancestral states. HM-INVs are ordered in columns, and each row corresponds to a species based on the phylogeny. Blue: the ancestral allele; red: the derived allele; gray: ambiguous state. The hierarchical clustering of HM-INVs and the lineage specificity annotation for HM-INVs are shown at the top. The log_10_-transformed length for each HM-INV is shown at the bottom, among which the HM-INVs with lengths > 95% quantile are indicated with red triangles. (**G**) The distribution of the normalized number of improperly aligned, paired-end reads for different groups of regions. M-INVs: macaque polymorphic inversions with high frequency; HS-INVs: human-specific inversions; Negative control 1 and Negative control 2: two groups of negative controls of shuffled regions (**Materials and Methods**). The median read number is shown above each boxplot. (**H**) UpsetR plot showing the number of human-specific inversions that are grouped based on their orientation relative to the reference genome across 18 haplotype assemblies. Blue: alignments concordant with hg38; red: inverted form; gray: ambiguous alignments. Human-specific inversions fixed in the population are highlighted with asterisks. Wilcoxon rank-sum tests, ***P value < 0.001. (**I**) Donut plot showing the classification of 101 candidate human-specific inversions based on their polymorphic states in human and macaque populations.

To verify the accuracy of the list of species-specific inversions between humans and macaques, we first compared it with a previous study reporting the species-specific inversions between humans and macaques based on Strand-seq data(4). 323 of the 375 inversions (86.1%) reported by Maggiolini *et al.* were also identified in this study, and we substantially expanded the list by including 1,649 additional inversions (**Fig. 4B**). Notably, the 1,649 additional inversions identified only in our study should also represent *bona fide*, species-specific inversion events. First, the list of species-specific inversions between humans and macaques was identified through comparative analyses of the assembled genomes, rather than *de novo* identification with short-or long-read sequencing data. In such a case, the boundaries of each species-specific inversion were supported by the original long reads used in the assembly of the human/macaque genomes, and the false positives possibly introduced by the segmental duplications nearby the inversions could be well controlled. Second, when performing pairwise genome alignments between human and other three old world monkeys, including crab-eating macaque, baboon and green monkey, 93.6% of these inversions were also supported by the genome assembly of at least one of these monkeys, providing additional verification of these events from the perspective of phylogenetic relationships. Third, we investigated why these 1,649 inversions detected in this study were not identified in the original Strand-seq data. To this end, we analyzed the 61 single cell libraries that were used to detect inversions in the original Strand-seq study. Notably, Strand-seq performs well to identify long inversions, while its sensitivity in identifying short inversions, or inversions with lower sequencing coverage, is relatively low. As expected, for the 1,649 inversions identified only in our study, when tracing the original Strand-seq signals, we found that the number of informative reads in these regions were actually significantly higher than that of the background (**Materials and Methods**, Wilcoxon rank-sum tests, P value < 2.2e-16; **Fig. 4C**), a pattern consistent with that of the shared list of 323 inversions (Wilcoxon rank-sum tests, P value < 2.2e-16; **Fig. 4C**). Moreover, for one discordant inversion between the two studies, we performed interphase fluorescence *in situ* hybridization (FISH, **Figs. 4D**, **4E** and **S8**, **Table S8**) with three position-specific oligo pools targeting the inversion region, which clearly supported the result in this study, in that the complex inversion was composed of two tandem inversions with breakpoint reuse, rather than a whole inversion as proposed by Maggiolini *et al*. (**Figs. 4D** and E)(4). This case study indicates that the method we proposed could even accurately clarify such complex, interlinked inversion events. Overall, the complete, accurate list of species-specific inversions between humans and macaques provides a basis for further delineating human-specific inversions.

We then performed phylogenetic analysis to trace the evolutionary trajectories of these species-specific inversions with the genome sequences from another nine non-human primate species (**Fig. 4F**, **Materials and Methods**). Notably, marmosets and squirrel monkeys were used as outgroups to infer the ancestral state of each inversion. In total, 1,240 of these inversions were adequately assigned the lineage information, including 101 human-specific inversions (**Table S9**), 48 macaque-specific inversions, 283 *Hominoidea*-specific inversions and 808 *Cercopithecidae*-specific inversions (**Fig. 4F**). Inversions with ambiguous ancestral states were excluded from the subsequent analyses. Notably, we covered 11 of the 12 human-specific inversions as previously reported(4) and included an additional 90 human-specific inversions. Interestingly, we found that inversions of *Hominoidea* origins (including human-specific and *Hominoidea*-specific inversions) were enriched with relatively larger inversions (Fisher’s exact test, P value = 0.046; **Fig. 4F**). Considering that the fixed inversions with larger sizes are likely to be maintained because of their adaptive functions, if these long inversions are fixed in humans, they may have contributed substantially to recent human evolution.

To further exclude polymorphic inversions not detected in the macaque reference genome but exist in macaque populations, we examined the states of the 101 candidate, human-specific inversions in the genomes of the 562 macaques. Briefly, the number of improperly aligned, paired-end reads in the region of interest was used as an indication of the existence of an inversion. Generally, in the homologous regions of these human-specific inversions in macaques, significantly fewer improperly aligned reads were identified, in comparison to polymorphic inversions with high frequencies in the macaque population (Wilcoxon rank-sum test, P value < 2.2e-16; **Fig. 4G**), indicating that the corresponding regions of most human-specific inversions were not polymorphic in the macaque population. Notably, we identified 13 deep polymorphic inversions that could be detected in both human and macaque populations (**Table S9**, **Materials and Methods**), which were removed from the list of candidate human-specific inversions.

Moreover, to further investigate whether the 88 human-specific inversions have been fixed in humans, we examined the state of each inversion in genome assemblies from 15 human individuals of multiple ethnicities (**Table S10**). Three human genomes with diploid assemblies were split into two haplotypes in this analysis. Notably, for 75 of the 88 human-specific inversions, the inverted state could be detected in all of these human genome assemblies (**Figs. 4H** and **I**), which were defined as fixed human-specific inversions (**Table S9**). Taken together, although we could not fully exclude the possibility that some of these inversions are still not completely fixed, according to their distributions in the 15 human individuals, it is more likely that they have been fixed.

### Human-specific inversions modulate the transcriptome in human brain

To study the features of these 75 fixed human-specific inversions, we first investigated whether they could introduce human-specific gene regulation through the reconfiguration of ancestral three-dimensional genome architecture. To this end, we first constructed Hi-C libraries from the prefrontal cortex (PFC) tissues of adult humans and macaques, and subsequently generated 1.3 billion valid contact pairs (**Table S11**). The quality of the Hi-C data was validated by distance-dependent interaction frequency decay and the ratio of *cis*-and *trans*-interactions (**Fig. S9A**, **Table S11**). Overall, a cross-species Hi-C map was developed to delineate hierarchical chromatin architecture, such as A/B compartments, TADs and loops (**Fig. S9**), with the map resolutions of 2.5 kb and 5.45 kb for human and macaque, respectively (**Table S11**).

We then compared the chromosome structures between these human-specific inversions and their orthologous regions in macaque, which was used as a proxy for determining their ancestral status, assuming that the chromosome structures of these regions have remained unchanged in the macaque lineage since its divergence from the human lineage. To investigate whether some of these human-specific inversions may be involved in the reconfiguration of gene regulation through chromosomal rearrangement after the divergence of humans and macaques, we identified inversions with coordinate overlapping with specific three-dimensional genome domains, such as compartments, TADs and loops. Interestingly, three of these inversions, associated with 47 protein-coding genes, are involved in the switching of TAD structures; and four human-specific inversions in the rewiring of preexisting chromatin loops, are putatively involved in the transcriptional regulation of nearby protein-coding genes. Among the four human-specific inversions associated with the chromatin loops, the largest one (chr18_inv1) introduces significant changes of the three-dimensional genomic architecture by disrupting the ancestral TAD domain and the chromatin loops around the breakpoints, which may further account for the differential expression of the associated genes such as *CLUL1*, *COLEC12* and *GREB1L* (**Fig. S10**). The other three inversions overlapped with the chromatin loops may have a similar association with the expression of the genes located in these regions.

Second, as previous findings suggest that the regions of inversion tend to accumulate mutations due to recombination suppression(25, 26, 28), we investigated whether these human-specific inversions could introduce regional mutation hotspots. Interestingly, we found that these regions harbored significantly more divergent sites than their adjacent regions in the human lineage since their divergence from chimpanzees (Wilcoxon signed rank test, P value = 1.3e-8 for the upstream regions, P value = 5.6e-9 for the downstream regions; **Figs. 5A** and **S11**). In contrast, in the orthologous regions of these human-specific inversions in macaques, the numbers of divergent sites were comparable with those in the adjacent regions (Wilcoxon signed rank test, P value = 0.56 for the upstream regions, P value = 0.68 for the downstream regions; **Fig. 5B**). Especially, we found an increased density of genetic divergence on promoter regions of genes located at these human-specific inversions, which may directly contribute to the cross-species differences of gene expression (**Fig. 5C**).

**Fig. 5.**
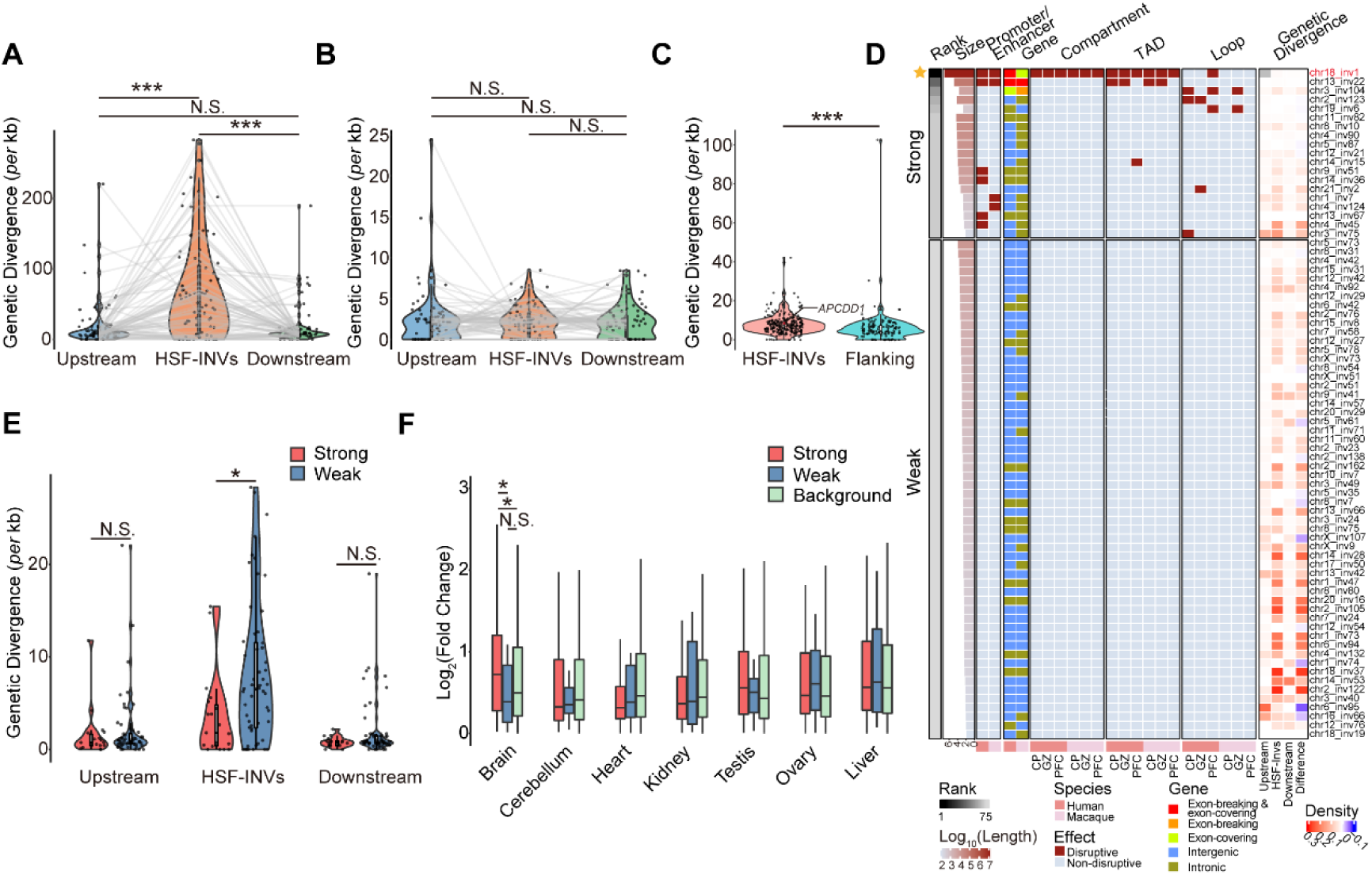
Characteristics of human-specific inversions. (**A**) Violin plots showing the genetic divergences of fixed human-specific inversions (**HSF-INVs**) and their length-matched, upstream and downstream genomic regions (**Upstream** and **Downstream** regions). Wilcoxon signed rank tests were performed. Wilcoxon signed rank tests, ***P value < 0.001. (**B**) Violin plots showing the genetic divergences of homologous macaque regions of the **HSF-INVs** and their upstream and downstream regions. Wilcoxon signed rank tests, N.S., not significant. (**C**) Violin plots showing the genetic divergences of promoter regions in fixed human-specific inversions (**HSF-INVs**) and promoter regions in length-matched flanking genomic regions (**Flanking regions**). The red dot indicates promoter of *APCDD1*. Wilcoxon rank-sum test, ***P value < 0.001. (**D**) Classification of 75 fixed human-specific inversions into two groups with different degree of regulatory effects (**Strong** and **Weak**), based on their sizes and locations in the human and macaque genomes. The genetic divergence relative to the human-chimpanzee common ancestor is also shown for the 75 **HSF-INVs** and corresponding upstream and downstream regions. The difference in the genetic divergence between each inversion and the average of its upstream and downstream regions is also shown (**Difference**). (**E**) Violin plots showing the genetic divergence for inversions with strong (**Strong**) or weak (**Weak**) effects, and their upstream and downstream regions. One-sided, Wilcoxon rank-sum tests, *P value < 0.05, N.S., not significant. (**F**) The log_2_-transformed fold changes in gene expression in the fetal brain between humans and macaques, for genes located on inversions with strong (**Strong**) or weak (**Weak**) effects, as well as for genome-wide orthologs as a background (**Background**). Wilcoxon rank-sum tests, *P value < 0.05, N.S., not significant.

As the polymorphic data within macaque population indicate that inversions with stronger effects are under stronger purifying selection, the inversions with stronger effects in humans (or not eliminated during the evolution) are thus more likely fixed due to their adaptive functions. Following this strategy, we then prioritized these human-specific inversions according to their sizes, effects on gene regulation and impacts on the three-dimensional genomic architecture (**Fig. 5D**). To this end, we ranked these human-specific inversions and classified them into two categories based on the strength of their effects (**Fig. 5D**; **Materials and Methods**). Interestingly, although these human-specific inversions generally showed higher level of divergence than the adjacent regions (**Fig. 5A**), the inversions with stronger effects showed a subtle increase in divergence, possibly due to a quicker pace of fixation shaped by positive selection (Wilcoxon rank-sum test, P value = 0.011; **Fig. 5E**).

Notably, for the fixed human-specific inversions with stronger effects, the resultant changes in transcriptome might be correlated with the direction of human adaptive evolution. Interestingly, the genes located around the inversions with stronger effects showed a greater degree of differential expression in fetal brain between humans and macaques, than genes associated with inversions with weaker effects (one-tailed Wilcoxon rank-sum test, P value = 0.028; **Materials and Methods**), or the genome-wide human-macaque orthologue pairs as a background (P value = 0.021), while no significant difference was detected in other tissues at different developmental stages, indicating the adaptive roles of these inversions in early brain development (**Figs. 5F** and **S12**). Our results thus provide a strategy for prioritizing human-specific inversions with adaptive functions in human evolution, especially in human brain development.

### A human-specific inversion contributes to unique human brain development

Among these 75 human-specific inversions, we focused on the inversion with the highest rank score (**Fig. 5D**, **Table S12**). This inversion was found to cover 13 Mb of chromosome 18, and included 62 protein-coding genes (**Figs. 5D** and **S10**). This human-specific inversion transferred the outward telomeric segment of the ancestral genome to the vicinity of the neocentromere in humans, and shifted the inward segment to the telomeric side. It disrupted the TAD structure near the inward breakpoint, and led to a loss of contact with previously adjacent region (**Figs. S10** and S**13**). Newly-formed interactions accompanying the inversion event were not detected, possibly due to the formation of a neocentromere in the human genome (**Figs. S10** and **S13**). Moreover, this inversion introduces mutation hotspots across the inversion region, which harbors more divergent sites than the adjacent region (density of divergent sites: 0.0073 *vs.* 0.0039 *per* bp). The reconfiguration of the genomic architecture and the introduction of the regional mutation hotspot should have contributed substantially to the human-specific changes of the transcriptome in this region: when comparing the expression levels of the human-macaque orthologues located near this inversion, we identified 43 differentially expressed genes between human and macaque, which is significantly higher than genomic background (Fisher’s exact test, P value = 0.033).

As a proof-of-concept, we focused on *APCDD1*, a gene located on this inversion and showed significantly-decreased expression in human brains in comparison to macaques and mice (Wilcoxon rank-sum test, P value = 1.6e-3; **Fig. 6A**). Consistent with the findings that inversions could introduce regional mutation hotspots, we found an increased density of genetic divergence on promoter regions of *APCDD1* in humans since the divergence from chimpanzees (**Fig. 5C**). We then designed dual-luciferase reporter assays to quantify the transcriptional activity of the *APCDD1* promoters in human and macaque, and found that the transcriptional activity of the *APCDD1* promoter sequence in humans is significantly lower than that in macaques (**Fig. 6B**, Student’s t test, P value < 1.0e-4). Next, we investigated whether the divergent sites accumulated during the evolution of the human lineage contributed directly to the changed activity. To this end, we identified five mutations occurred specifically in humans after the divergence of human and chimpanzee, which are located on *APCDD1* promoter regions and marked by H3K4me3 signals. When we mutant these sites to their ancestral alleles, three of them could significantly enhance the transcriptional activity of the promoter (**Fig. 6B**, Student’s t test, Mutation-1: P value < 1.0e-4, Mutation-3: P value = 1.0e-4, Mutation-5: P value = 1.6e-3). These findings thus support that the accumulated mutations on *APCDD1* promoter, a process accelerated by the formation of this human-specific inversion, contributed to the decreased *APCDD1* expression in humans.

**Fig. 6.**
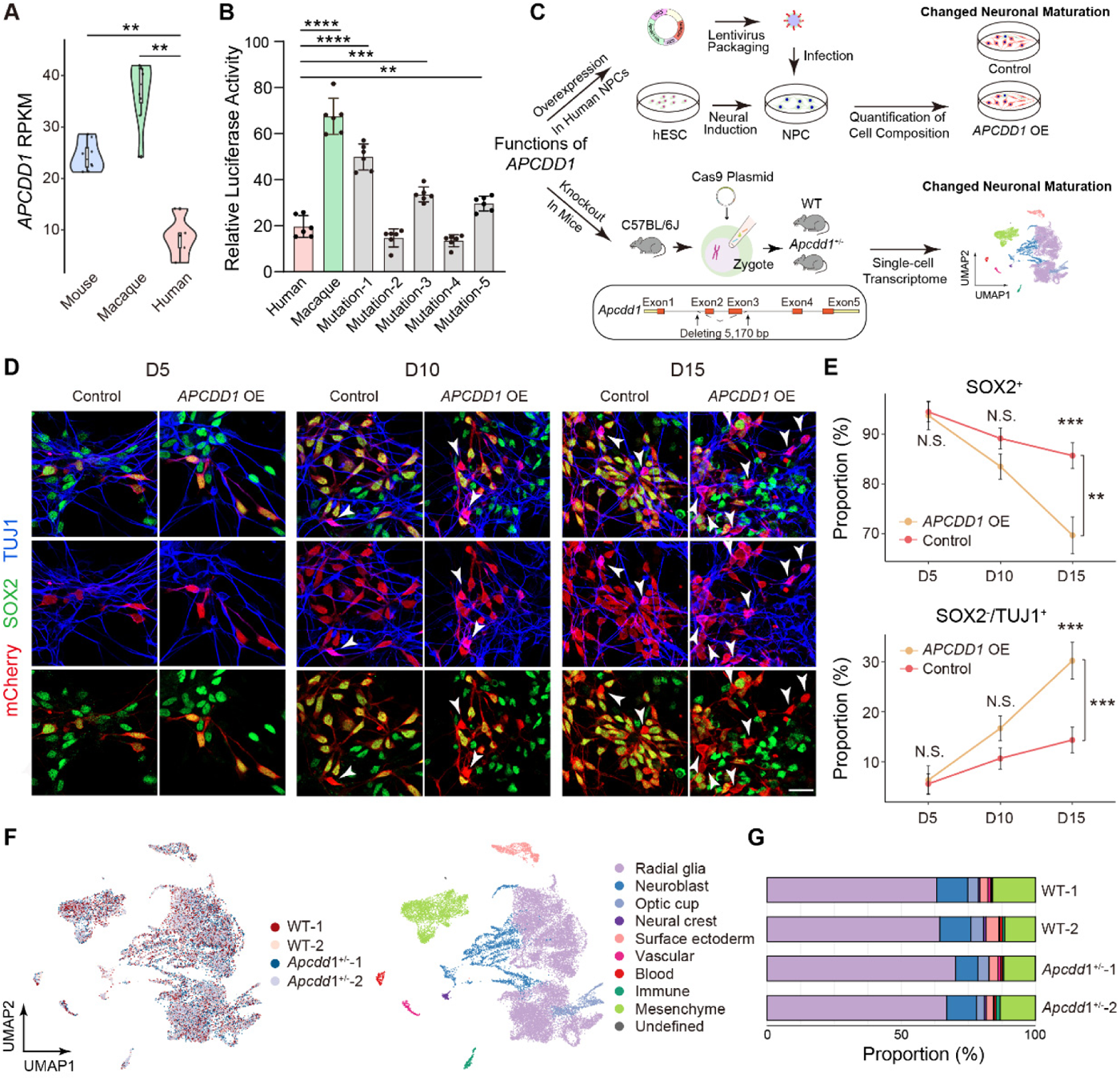
A human-specific inversion contributes to human brain development. (**A**) Violin plots showing the expression levels of *APCDD1* in the brains of humans, macaques and mice at the mid-to late-fetal stages. Wilcoxon rank-sum tests, **P value < 0.01. (**B**) Relative luciferase activities (*Firefly*/*Renilla*) of *APCDD1* promoter in human (**Human**), *APCDD1* promoter in rhesus macaque (**Macaque**), and *APCDD1* promoter in human with five mutations **(Mutation-1**, **Mutation-2**, **Mutation-3**, **Mutation-4**, **Mutation-5**). Student’s t test, **P value < 0.01, ***P value < 0.001, ****P value < 0.0001. (**C**) Experimental design for *APCDD1* functions. The experimental design for functional studies of *APCDD1*, including the *APCDD1* overexpression assay in NPCs and the functional studies in *Apcdd1*+/-mice. (**D**) Representative immunostaining of SOX2 (indicative of progenitors), TUJ1 (indicative of postmitotic neurons) and mCherry (indicative of virus-infected cells) in the assays of wild-type NPCs (Control) and NPCs with *APCDD1* overexpression (*APCDD1* OE), on protocol day 5 (D5), day 10 (D10) and day 15 (D15) after lentiviral infection. The white arrows indicate SOX2^-^/TUJ1^+^ cells. Scale bars, 20 μm. (**E**) Proportions of SOX2^+^ progenitors (top) and SOX2^-^/TUJ1^+^ neurons (bottom) in the Control and *APCDD1* OE assays on different protocol days (D5, D10 and D15). Data are represented as the means ± SEMs. Two-way ANOVA, **P value < 0.01, ***P value < 0.001. (**F**) UMAP plots of scRNA-seq from brains of two replicates of wild-type (**WT-1** and **WT-2**) and *Apcdd1^+/-^* mice (***Apcdd1^+/-^*-1** and ***Apcdd1^+/-^*-2**) at E10.5, grouped by genotypes (left) and cell types (right). (**G**) Proportion of each cell type in brains of **WT-1**, **WT-2, *Apcdd1^+/-^*-1** and ***Apcdd1^+/-^*-2**.

To investigate whether the decreased expression of *APCDD1* is involved in the adaptive evolution of human brain development (**Fig. 6C**), we first developed a lentiviral-based overexpression assay in human neural progenitor cells (NPCs) to examine the effects of *APCDD1* overexpression on the proliferation and differentiation of neural progenitors (**Figs. 6D** and **E**). Notably, we found a significant reduction of SOX2^+^ progenitor populations (two-way ANOVA and post hoc Tukey’s HSD test, P value = 1.1e-3) and an expansion of SOX2^-^/TUJ1^+^ neuron populations (P value = 9.9e-4; **Figs. 6D** and **E**), indicating an accelerated neuronal maturation process in human NPCs with *APCDD1* overexpression, a pattern recapitulating the differences in early brain development between humans and macaques(30).

As the varied pace of neuronal maturation has been implicated into the different features of brain development(31), we further constructed *Apcdd1^+/-^* mice based on CRISPR/Cas9 technology to investigate whether the partial depletion of *Apcdd1* could contribute directly to the alteration of cell composition in the mouse brain, a distinct feature arising during human adaptive evolution (**Figs. 6C**, **6F**, **6G** and **S14**). To this end, we performed single-cell RNA sequencing (scRNA-seq) in the brains of wild-type and *Apcdd1^+/-^* mice at E10.5 and obtained a transcriptional profile of a total of 25,074 cells (**Materials and Methods**). The scRNA-seq data from each genotype were then integrated to determine the cell type diversity in mouse brains. Cell type identities were assigned based on known marker genes as reported in literatures(32–36), which resulted in nine major classes including radial glia, neuroblast, optic cup, neural crest, surface ectoderm, vascular, blood, immune and mesenchyme (**Figs. 6F** and **S15**). Notably, in contrast to wild-type mice, we found an increased proportion of radial glial cells in *Apcdd1^+/-^* mice (**Fig. 6G**), recapitulating the involvement of *APCDD1* in the regulation of neurogenesis in human NPCs.

These findings jointly indicate the regulatory effect of *APCDD1*, possibly through a human-specific inversion, on neuronal maturation that are distinct in human.

## Discussion

In this study, to improve the variant calling in macaque populations, we first attempted to construct a better reference macaque genome, through the integration of three recent genome assemblies based on third-generation sequencing. Notably, the strategy of gap filling by aligning Mmul_10 to the other genome assemblies may introduce assembly errors of other assemblies into Mmul_10. To exclude this possibility, we used a very conservative strategy in this step, in which only gaps with continuous sequences spanning the gap regions were filled (**Materials and Methods**). Moreover, we further sequenced the genome of eight additional macaques with PacBio long HiFi reads, and assessed these regions of gap filling with these HiFi reads. According to the alignments of these long reads on the Mmul_10 genome assembly with gap closures, we found that all of the 40 gap closures could be validated by these long reads (**Fig. S2**), indicating that these gaps should represent real genomic gap regions of Mmul_10, rather than the assembly errors from other macaque genome assemblies, or individual-specific structural variations.

SVs, with larger sizes and expected stronger effects than SNVs, have been reported as one of the major sources of genomic innovations. However, it is error-prone to use standardized algorithms to call SVs with short reads. Notably, despite the inherent advantage of long-read sequencing in pinpointing SVs in a straightforward manner, the cost of this strategy limits its application at a large population scale, and the SVs identified by long-read data in a small number of macaques could only provide limited vision for the complete SV atlas in macaque population, hindering the in-depth investigations of their global features and evolution. A systematic workflow for reliable SV identification with abundant short-read sequencing data is thus urgently needed. Here, we used an optimized pipeline for accurate SV detection with short reads, in which different SV tools were integrated and cross-validated to reduce false positives. With this new pipeline, we present the largest macaque SV landscape to date, based on the deep integration of WGS data of 562 macaques. The high verification rate as indicated by the evaluations with public SV benchmarks, an inhouse benchmark of eight macaque genomes with long-read sequencing, and another benchmark of one macaque genome with long-read sequencing and whole genome assembly, jointly verified the efficiency of our pipeline in accurately calling SVs in macaque populations. This accurate and quantitative SV map thus provides a basis for clarifying the features, turnover and evolutionary significance of SVs in primates.

Although it is difficult to accurately identify inversions with short reads, according to our evaluations with the three-tier benchmarks, our pipeline actually performs well in calling inversions, with the verification rate comparable with deletions. Moreover, we found stronger signals of purifying selection on the fixation of inversions than other types of SVs, consistent with previous reports that polymorphic inversions are largely deleterious due to recombination suppression and the subsequent accumulation of deleterious mutations. We thus focused on this type of SVs with possibly stronger effects during the human evolution in this study.

Based on the macaque SV map and a comparative genomics study with multiple outgroup species, we identified a list of 75 human-specific inversions apparently fixed in humans. Moreover, using our quantitative SV map in macaque population, we found that the inversions located in regulatory regions, such as genic regions and TAD boundaries, are subjected to enhanced purifying selection, suggesting a practical strategy for prioritizing these human-specific SVs with the most important functions in shaping human adaptive evolution. Accordingly, when we classified these human-specific inversions into two categories on the basis of the strength of their effects, we found that the genes located in inversions with stronger effects showed a higher degree of differential expression in fetal brains between humans and macaques, which was well correlated with the direction of human adaptive evolution in brain development. These fixed, human-specific inversions with stronger effects should thus have substantially shaped the human brain transcriptome during human evolution, through their dual effects of reconfiguring ancestral genomic architecture and introducing regional mutation hotspots.

Our findings suggest that the accumulated mutations on *APCDD1* promoter, a process accelerated by the formation of this human-specific inversion, should have contributed to the decreased expression in humans. However, the reconfiguration of the ancestral chromosome structure may also be involved in this regulation. Notably, along with the emergence of this human-specific inversion, the locus of *APCDD1* was moved to a region near the newly-formed centromere in human. Further investigations are thus needed to justify the effects of changed chromosome structure on *APCDD1* expression during the human evolution. The contribution of regulatory changes to human evolution has been proposed to supplement contribution of sequence alterations in coding regions. The proof-of-concept study of *APCDD1* regulation indicates that the human-specific downregulation of even one single gene, presumably regulated by a human-specific inversion, could significantly modulate the process of neuronal maturation, recapitulating the unique features of human brain development. The human-specific inversions identified here could thus act as a previously neglected genetic source underlying the uniqueness of human brain development.

## Materials and Methods

### Ethics statement

The rhesus macaque samples used in this study were approved by the Animal Care and Use Committee of Peking University (IMM-LiCY-1). The human PFC sample in this study was approved by the IRB of Beijing Tiantan Hospital, Capital Medical University (KY2017-035-02).

### Sample collection

#### Human brain tissue

Human brain tissue was obtained from Beijing Tiantan Hospital.

#### Rhesus macaque brain tissue

Frozen brain tissues of 26 rhesus macaques were obtained from the animal facility at the Institute of Molecular Medicine, Peking University (accredited by the Association for Assessment and Accreditation of Laboratory Animal Care, AAALAC). Detailed information for these macaques is provided in **Table S4**.

#### HeLa-S3 cell line

The HeLa-S3 cell line was obtained from the laboratory of Dr. Yujie Sun, Peking University. HeLa-S3 cells were grown in 10 mm glass-bottom imaging dishes (Cellvis, #D35-10-1-N) with 2 mL of modified medium (high-glucose DMEM, Thermo Fisher Gibco, #10569-044) supplemented with 10% (v/v) fetal bovine serum (Thermo Fisher Gibco, #10091-148) and 100 U/mL penicillin‒streptomycin antibiotics (Thermo Fisher Gibco, #15070-063) under regular cell culture conditions (37 °C, 5% CO_2_, humidified atmosphere). The cells were passaged at a proportion of 1:8-1:10 with trypsin (Life Technologies, #25200-056) every 3 days or when they reached 80% confluence.

#### LLC-MK2 cell line

Macaque Lilly Laboratories Cell-Monkey Kidney 2 (LLC-MK2) cells were obtained from the National Collection of Authenticated Cell Cultures (CSTR: 19375.09.3101MONGNO6). The LLC-MK2 cells were maintained *in vitro* in RPMI 1640 medium (Thermo Fisher Gibco, #C11875500BT) supplemented with 10% (v/v) fetal bovine serum (Thermo Fisher Gibco, #10091-148) and 100 U/mL penicillin‒streptomycin antibiotics (Thermo Fisher Gibco, #15070-063) under cell culture conditions (37 °C, 5% CO_2_, humidified atmosphere). The cells were passaged at a proportion of 1:3-1:4 every 3 days.

### Deep sequencing

Genomic DNA was extracted from frozen brain samples using Ultra Pure^TM^ Phenol:Chloform:Isoamyl Alcohol (Invitrogen, 25:24:1, v/v). For PacBio sequencing, SMRTbell genomic libraries (> 20 kb in length) were constructed from high-quality DNA extracted from the brain and muscle tissues of one male macaque, and deep sequencing was performed on the RS II and Sequel platforms with the P6-C4 sequencing reagent. For whole-genome sequencing with short reads, libraries were constructed from DNA extracted from brain or blood tissues of 27 macaques, and deep sequencing was performed on MGISEQ-2000 (MGI) sequencing systems to generate 150 bp, paired-end reads.

### Hi-C library preparation

Frozen PFC samples (0.2 g) from a human and a macaque were ground in liquid nitrogen, after which cell suspensions were prepared. The filtered cell suspensions were fixed in 1% formaldehyde for 30 min. The cross-linked DNA was digested with 200 U MboI (NEB, #R0147M), and the digested fragment ends were filled with biotin-14-dATP (Invitrogen, #19524016), dCTP, dGTP and dTTP by DNA Polymerase I Klenow Fragment (NEB, #M0210L). The resulting blunt-end fragments were religated by T4 DNA ligase (Enzymatics, #L603-HC-L). Then, the ligated cross-linked DNA was reversed using proteinase K (TIANGEN, #RT403). DNA purification was performed by removing biotin from unligated ends using T4 DNA polymerase (Enzymatics, #P7080L). The purified DNA was then sheared to a length of 350-500 bp using Covaris M220, and biotin-labeled DNA was pulled down with Dynabeads M-280 Streptavidin (Invitrogen, #11205D). Library preparation was performed using a DNA library preparation kit (Vazyme, #ND607). The libraries were then sequenced on the Illumina NovaSeq 6000 sequencing platform to generate 150 bp paired-end reads.

### Gap closure

Gap closure was performed according to the genome assembly of Mmul_10. Genome-wide pairwise alignment was performed using lastz (Parameters: --notransition --step=20 -- ambiguous=n --chain)(37) between Mmul_10 and two other reference genomes, Mmul_8.0.1 (Indian-origin) and rheMacS (Chinese-origin). A gap was closed when two conditions were met: 1) there was a continuous sequence spanning the gap region in alignment with Mmul_8.0.1 or rheMacS, and 2) the flanking anchor sequences along the gap regions were longer than the gaps. The sequences spanning the gap regions were extracted as padding sequences. If a gap in Mmul_10 could be filled based on both of the genome assemblies, the rheMacS genome was preferentially used, considering its higher genome continuity.

To evaluate the quality of gap closure, a Bionano optical map was generated for one macaque (monkey ID: blood-270-zhongxm_HKNGYCCXX_L1). Briefly, to create a Bionano optical map, DNA was extracted from the liver of the same macaque used for PacBio sequencing and digested with the nicking enzyme NT.BspQI. Standard library preparation and optical scanning were then performed. Bionano Solve software was used to assemble the optical map from scratch to obtain scaffolds (version: 3.1), and then the script fa2cmap_multi_color.pl was used to obtain the optical map scaffold with enzyme digestion (enzyme name: BspQ). OMTools (version: 1.4a)(38) was used for read mapping and visualization. Only gap sequences with at least one mapped restriction enzyme cleavage site or with a consistent order of restriction enzyme cleavage sites around the gap were retained.

Ten macaque samples with an average sequencing depth of 36 fold (5 Chinese-origin macaques and 5 Indian-origin macaques) were selected to evaluate the quality of gap closure. The whole-genome sequencing data of these samples were aligned to the improved genome assembly with 44 gaps filled using BWA-MEM(39) with the default parameters (version: 0.7.17-r1188). Then, the read coverages of the gap regions and their length-matched, upstream or downstream regions were calculated and compared using bamdst with the default parameters (version: 1.0.9).

Another eight Chinese-origin macaques with PacBio long-read sequencing data were further selected to assess these regions of gap closure (**Table S3**). The long reads of these samples were aligned to the improved genome assembly with 40 gaps filled, using pbmm2 with default parameters (version: 1.8.0). The alignments of these regions were then illustrated using IGV(40) with default parameters (version: 2.16.0).

### Identification and annotation of SNVs in the macaque population

The whole-genome sequencing data of rhesus macaques were aligned to the genome assembly of rheMac10Plus with BWA-MEM using the default parameters (version: 0.7.17-r1188). PCR duplicates were marked for the merged BAM files using GATK MarkDuplicates (version: 4.2.2.0), and only non-duplicate reads were used for the downstream analyses.

The best-practice workflows of the Genome Analysis Toolkit (GATK, version: 4.2.2.0) were used to call SNVs. Indels were realigned with IndelRealignment, and base quality was recalibrated with BaseRecalibrator. The SNVs on each chromosome in each sample were called with HaplotypeCaller. The known SNV datasets were obtained from recent macaque population studies(16, 19), which were used to evaluate the variants with VariantRecalibrator, with a threshold of sensitivity above 99%. To ensure the accuracy of variants, the evaluated variants were then used as the new “known SNV dataset” to call and evaluate the SNVs as illustrated in the above workflow. The PLINK tool (version: v1.90b6.16 64-bit)(41, 42) was further used to filter the samples with frequencies of missing calls greater than 10% and low-quality SNVs with the following exclusion criteria: 1) variants with frequencies of missing calls greater than 10%, 2) variants with Hardy-Weinberg equilibrium exact test P values below 0.001. The SnpEff tool (version: 5.0e)(43) was then used to annotate these SNVs.

### Kinship analyses

The relationships between individual macaques were estimated using KING software (version: 2.2.5)(44). The kinship coefficients for all pairs of macaques were estimated and assigned to one of the following categories: twins, parent-offspring, siblings, 2nd-degree relatives or 3rd-degree relatives, using the recommended kinship coefficient boundaries(44). By considering the family trees, we filtered coefficients for pairs of relatives inferred to be 2nd-degree relatives or closer to identify a list of independent macaque animals(45).

### Population genetic analyses

For both human and macaque populations, the nucleotide diversity (π) was calculated for genome-wide sliding windows (3 Mb for each window and 3 Mb for each step) using VCFtools (version: 0.1.17)(46).

### Evaluation of the reference-bias in reads mapping

We first identified the inherent divergent regions between Indian-origin and Chinese-origin populations of rhesus macaques, by aligning the Indian-origin reference genome (rheMac10Plus) with the Chinese-origin reference genome rheMacS, and with the inhouse-assembled genome of a Chinese-origin macaque. The inherent divergent regions were then defined as the overlap of divergent regions of the two genome-wide alignments. Notably, regions of 40 filled gaps were not considered in the definition. We then identified SNVs located on these regions, and analyzed the frequencies of the derived allele of these SNVs in different subpopulations.

### Genetic diversity and population structure analyses

A neighbor-joining phylogeny for 144 Chinese-origin macaques was constructed based on the P distance matrix as calculated by VCF2Dis (version: 1.47), according to the autosomal genetic variations. The FastME tool (version: 2.0)(47) was used to visualize the phylogenetic tree. The subpopulation information was assigned for animals with an unknown geographic distribution on the basis of the clustering results and the genetic distance. Subpopulations of *M. m. mulatta* and *M. m. lasiotis* were combined due to their admixture structures. PCA of SNVs on all of the autosomes in the macaque population was performed using GCTA (version: 1.93.0)(48). A PCA scatter plot for principal components 1-3 was then created using the plotly package in R (version: 4.1.2). The fractions of the variance explained by the three components were calculated according to the Kaiser-Guttman criterion and the broken-stick model.

### Identification of SVs in the macaque population

PCR duplicates were marked for the raw mapped reads using Picard (version: 2.25.4) and then structural variants of each animal were identified using DELLY (version: 0.8.2)(49) and SpeedSeq (version: 0.1.2)(50). Multiple filters and intersection strategies were used to obtain an accurate set based on the evaluation of macaques with genome assemblies. Specifically, only SVs with “QUAL=PASS” were retained from the DELLY results. Among all SV calls from DELLY and SpeedSeq, only SVs with lengths of at least 50 bp on autosomal and X chromosomes were retained. For the results of CNVnator, thresholds were set as follows: 1) both E-val 1 and E-val 2 were less than 0.05; 2) q0 was less than 0.5 and not equal to -1.

Inversions, deletions and duplications were then identified separately. Briefly, for candidate inversions, only SpeedSeq calls supported by at least three paired-end reads and three split reads were retained. In addition, only inversions with a consensus region shared by DELLY and SpeedSeq calls (with a coordinate overlap of at least 90%) were retained. For candidate deletions, only DELLY calls supported by at least three paired-end reads and three split reads were retained. Moreover, only deletions with a consensus region shared by DELLY and SpeedSeq calls (with a coordinate overlap of at least 50%) were retained. Notably, we removed the deletion calls showing at least 20% overlap with gap regions in the macaque reference genome and retained only candidate deletions showing significantly lower read depths in the deletion regions in contrast to their adjacent regions. For candidate duplications, only DELLY calls supported by at least one split read were retained. Moreover, only duplications with a consensus region shared by DELLY and SpeedSeq calls (with a coordinate overlap of at least 50%) were retained. Candidate duplications located in repeat regions were retained in the following analyses only when they were also identified by the CNVnator tool (with a coordinate overlap of at least 80%).

For each SV type, the SV calls from the macaque population were then merged into a single callset with the svtools tool (version: 0.0.1)(51), which was used as a reference to genotype each macaque animal. Considering the limited resolution in identifying breakpoints, we clustered and merged the intersecting SV calls to obtain consensus SV hotspots with BEDtools merge. Overlapped SV calls were merged into a single interval. Per-sample quality control was then performed according to these SV hotspots. Notably, we removed 10 macaque samples in which the SV counts were >10 median absolute deviations (MAD) from the median count of the corresponding SV type. Allele frequencies were then recalculated based on the SV hotspots. The genotypes were assigned as “missing” for individual samples if there were ambiguous genotypes on this SV. Samplot (version: 1.3.0)(52) was used to visualize the SV calls.

The SV calls in individual macaques were validated with both long-read sequencing data and an intensive manual check. Vapor (version: 1.0)(53) was used to validate the SV calls from the macaque with long sequencing reads. The ambiguous calls were visualized with the SV-plaudit tool (version: 2.0.0)(54) for a manual check.

### Development of a three-tier benchmark

We first evaluated the performance of our pipeline in SV detection with public benchmarks. Briefly, we selected the HG002 genome with verified deletion calls as the benchmark callset(21), and evaluated the performance of our methods in calling these deletion events, by processing the short-read sequencing data of this genome (SRR12898337, data generated by Illumina HiSeq X Ten with a coverage of 42 fold), and benchmarking using truvari tool provided by the GIAB(55). We selected the SV calls with SVTYPE=DEL and performed truvari bench with default parameters (--pctsim=0: to ignore sequence comparison, --includebed HG002_SVs_Tier1_v0.6.bed file: to benchmark in high-confidence regions, --passonly: to filter out the highest confidence set of SVs). As this callset does not include inversion events, we then used another benchmark callset from Pendleton *et. al*(22) to evaluate the performance of our pipeline in calling inversions, by processing the short-read sequencing data of the HG001 genome (SRR8454588, data generated by Illumina HiSeq 4000 with a coverage of 30 fold), and benchmarking using BEDtools tool.

Second, from the list of 562 macaques, we selected eight macaques and sequenced their genomes with different coverages of long HiFi reads (**Tables S3**). For each macaque animal, we then evaluated the performance of our pipeline in calling deletions, duplications and inversions, using SVs called in this macaque by long reads as a benchmark. As the sequencing of these samples was not saturated, for each SV type in each macaque animal, we performed a simulation step to estimate the theoretical number of verified SVs at the current sequencing depth, assuming the SVs and their genotypes were accurately identified with short reads. The verification rate was then calculated by dividing the detected number of verified SVs by the mean of the theoretical number of verified SVs from 10,000 times of simulations. If the calculated verification rate exceeds 100%, it was regarded as 100%.

Finally, from the population of 562 macaques, we selected the macaque with short-read sequencing data and the Bionano optical map generation (as mentioned in the section of “gap closure”), and further sequenced its genome with high-coverage, long-read sequencing. We further *de novo* assembled its genome on the basis of the integration of the short-read sequencing, long-read sequencing and Bionano optical data. Briefly, long-read *de novo* genome assembly was performed with FALCON software. Contigs were further polished with Arrow using long-read sequencing data, and Pilon(56) (version 1.2) using short-read sequencing data. *De novo* genome assembly with Bionano optical map data was performed by BioNano_Solve (version Solve3.1_08232017) with manufacturer recommended parameters. Hybrid scaffolding of the polished PacBio contigs and Bionano optical map were performed by the hybrid scaffolding module packaged with the BioNano_Solve software. The genome assembly of rheMac8 assembly and the genomic markers were then used to generate chromosome-level assembly by ordering the hybrid scaffolds and contigs. Finally, the ordered hybrid scaffolds and contigs were linked together by filling with 3 Mbp N sequences between adjacent scaffolds or contigs. The long reads and the de novo assembled genome were then used as the benchmark to evaluate the performance of our pipeline in SV calling with short reads.

### PCA of SVs

PCA was performed with the R command prcomp based on the genotype information for 562 unrelated animals, in which the macaques were classified into three groups: captive Indian-origin macaques, captive Chinese-origin macaques and wild-caught Chinese-origin macaques.

### Hi-C data analyses

For the Hi-C data of the adult PFC and fetal brain (CP and GZ zones) tissues of humans and macaques, we first performed preprocessing with HiC-Pro (version: 2.11.1)(57). Paired-end reads in FASTQ format were first aligned against the reference genome (hg38 for human and rheMac10Plus for macaque) and were subjected to rigorous filtering criteria to generate valid contact pairs. We then calculated the resolution of the maps as described in Rao *et al.*(58). The valid pairs were then binned to contact maps at different resolutions, which were subjected to ICE normalization. The Juicer toolkit (version: 1.9.9)(59) was used to convert the contact maps to .hic files for visualization in the UCSC Genome Browser. Knight-Ruiz (KR) normalization is used for configuring Hi-C tracks. The analysis of distance-dependent contact strength decay was performed using FAN-C1 (version: 0.9.21)(60). We then detected basic chromosome organizations in a hierarchical manner. First, Cooltools (version: 0.3.2)(37) was used to identify A/B compartments at a 100 kb resolution, with the genome divided into two distinct groups based on the sign of the first PC (PC1). The group with higher gene density was defined as compartment A, and the other was compartment B. Next, TAD boundaries were detected using the insulation score method at a resolution of 40 kb. Consequently, TAD bodies were defined as genomic regions between two adjacent boundaries. Finally, chromatin loops were called by HiCCUPS (version: 1.9.9)(58) at 10 kb and 25 kb resolutions, and significant interaction pairs were detected using HOMER (version: 4.1.1)(61) at 25 kb resolution. Aggregated peak analysis (APA) was performed using the Juicer toolkit to measure the enrichment of the detected chromatin loops. Notably, as a consequence of the discrepancy in the sequencing depth achieved for adult PFC tissues between humans and macaques (approximately 3:1), we downsampled the contact pairs for human data in proportion to the genome assembly size in the cross-species comparisons.

### Prediction of promoter and enhancer regions in the macaque brain

ChIP-seq data of eight distinct anatomical regions of the macaque brain were downloaded from GEO (GSE67978)(62). Sequencing reads were aligned to the rheMac10Plus reference genome with Bowtie (version: 1.3.0)(63). Peak calling was performed with MACS2 (version: 2.2.7.1, parameters: P = 10^−5^ extsize = 400 local lambda = 100,000)(64) for the ChIP-seq data of H3K27ac and H3K4me3 marks. To match the peak resolution of histone marks, peaks smaller than 2,000 bp were extended to 2,000 bp(65, 66). For H3K27ac modification, the enriched regions identified in at least two biological replicates were defined as reproducible enriched regions. For H3K4me3, as only one biological sample was sequenced, all peaks called were retained. The regions marked by peaks of both H3K27ac and H3K4me3 and located within 1,000 bp around the transcription start sites (TSSs) were defined as putative promoter regions. The regions marked by peaks of H3K27ac and located 1,000 bp away from TSSs were defined as putative enhancer regions. The peaks were annotated with ChIPseeker tool (version: 1.22.1)(67).The final list of predicted promoter and enhancer regions in the macaque brain was obtained by merging the identifications from all of these brain samples.

### Inference of the ancestral state for inversions and deletions in the macaque population

Genome assemblies of human (hg38), marmoset (calJac4) and rhesus macaque (rheMac10Plus) were used to trace the ancestral state of inversions and deletions in the macaque population. For inversions, we downloaded the chain/net alignment across humans and marmosets/macaques from the UCSC Genome Browser. The regions labeled with “INV” in the net alignment file were extracted as potential inverted regions. The inversions called within the macaque population were then compared with the annotations in these species, and the ancestral state of each inversion was then defined. Inversions with no authentic alignments or ambiguous ancestral states were excluded from the subsequent analyses. For deletions, we manually generated multiple alignment format (MAF) files between rhesus macaques and humans or marmosets using netToAxt. To decide whether each deletion variant represented a deletion within the population or an insertion in the reference genome, we then partitioned the genomic context of each deletion into two parts: the deletion region and the anchor regions (the upstream and downstream 200 bp regions). Basewise alignments of these two parts were extracted from the MAF files, and the alignment state was then determined if the alignments in both upstream and downstream regions could be anchored. The state (deletion or not) was then determined based on the alignment length of the deletion regions. The ancestral state was thus defined according to the alignments in these species. Deletions with no anchor alignments, inaccurate alignment lengths or ambiguous ancestral states were excluded from the subsequent analyses.

### Classification of population inversions based on regulatory regions

For the annotations of inversions from the perspective of genome architecture, we analyzed public Hi-C data from the fetal brain (CP and GZ zones)(29) and defined TAD boundaries as the 10 kb intervals (± 5 kb) centered on the start and end coordinates of each TAD. Inversions showing overlap with TAD boundaries were defined as inversions disrupting TAD boundaries. For the annotation of inversions according to gene structures or functional regions, we extracted the gene structure annotations from the rheMac10Plus assembly and defined the sequences outside of all annotated genes as intergenic regions. Promoter and enhancer regions were predicted from the ChIP-seq data of macaque brains. Inversions overlapping with exons were defined as inversions disrupting exonic regions. Inversions overlapping with predicted promoter and enhancer regions were defined as inversions disrupting functional regions.

### Detection of inversions between humans and macaques

We used the DoBlastzChainNet.pl script deposited in Kent’s utilities of the UCSC Genome Bioinformatics Group to construct a chain/net alignment across the human (GRCh38/hg38) and macaque (rheMac10Plus) genomes. We first set up the alignment with a chain/net of hg38 as the target and rheMac10Plus as the query (-chainMinScore=5000 – chainLinearGap=medium -syntenicNet), and the alignment in the reverse direction was then generated in “swap” mode (-swap -syntenicNet). The main method implemented with this script was pairwise whole-genome alignment using lastz (version: 1.04.00), with primate-specific parameters specified, which can be found at http://genomewiki.ucsc.edu/index.php/DoBlastzChainNet.pl#lastz_parameter_file. Then, the genomic regions that were labeled “INV” in the net alignment file and reciprocally intersected in both directions were extracted as candidate inversions, and those with over 80% of the regions falling within SDs, as annotated in hg38, were filtered out. We further performed manual curation of each net inversion by revising the breakpoints of inversions, rescuing nested inversions showing breakpoint reuse, and removing ambiguous cases. Finally, 1,972 species-specific inversions across humans and rhesus macaques were retained (**Table S7**).

### Validation of species-specific inversions between humans and macaques

#### Comparison with Strand-seq inversions

As strand-seq studies typically report candidate inversions with broad boundaries, we used a relatively loose threshold (more than 10% coordinate intersection in the two studies) in determining the overlaps, which were then subjected to a manual check on the UCSC Genome Browser to confirm that these inversions were indeed identified by both methods. Notably, when calculating the overlap for the body regions of these inversions identified separately by the two methods, the mean percentage of the overlapped regions reaches up to 95.8%.

#### Validations with phylogenetic relationship

We performed pairwise genome alignments between human and other three old world monkeys, including crab-eating macaque, baboon, and green monkey, and checked whether these inversions were supported by the genome assembly of these closely-related monkeys.

#### Strand-seq validation

The bam files for 61 single-cell Strand-seq libraries derived from one macaque lymphoblastoid cell line (LCL) were downloaded from NCBI BioProject under accession number PRJNA625922. We discarded supplementary alignments and only retained properly-paired reads. To validate inversions with Strand-seq data, we first defined the template states (CC: plus-plus-strand, WW: minus-minus-strand or WC: plus-minus-strand) for the flanking regions of these inversions, as described in Porubsky *et al.*(*68*). If the template states of the upstream and the downstream regions of the candidate inversion were identical, then the reads in the opposite direction in the inversion body region were defined as informative reads. Notably, since the removal of newly synthesized strand may not be perfect, low abundance reads in the opposite direction are expected even for WW or CC chromosomes, typically at an average noise level of 5%. Accordingly, we estimated the background for the number of falsely assigned informative reads by assuming a 5% error rate probability. Only the cases where the reads coverage ≥ 3 at the inversion loci in the Strand-seq data were included in the evaluations.

#### FISH validation

To determine the state of discordant inversions between this study and a previous report(4), we chose three conserved genomic regions in human and macaque genomes to design probes. Through the order of these distinct probes in the FISH assay, we could distinguish between the two models (**Fig. 4D**). Oligonucleotide probes were designed using Oligominer (version: 1.7) following the online instructions and the method described in Boettiger *et al.*(69). The template oligo pools designed for all six genomic regions were synthesized by Synbio Technologies (Suzhou, China), and the flanking primer binding sequences used to amplify the probes are shown in **Table S8**. The probes were amplified from the oligo pools as described in Li *et al.*(70). For the FISH experiments, HeLa-S3 cells and macaque LLC-MK2 cells were first synchronized in metaphase through 16 h of nocodazole treatment, with final concentrations of 50 ng/ml and 20 ng/ml, respectively. Subsequently, cells collected under trypsin (Life Technologies #25200-056) treatment were exposed to a hypotonic solution (10 mM KCl for HeLa-S3 cells, 30 mM KCl for macaque LLC-MK2 cells) for 15 min and fixed in 3:1 methanol:acetic acid. Metaphase spreads were then prepared and stained with DAPI according to the methods of Li *et al.*(71). Finally, FISH experiments were conducted, and imaging was performed on a Nikon Live SR CSU-W1 Spinning Disk Confocal microscope as described in Li *et al.*(70).

### Identification of human-specific inversions

To trace the evolutionary trajectory of each species-specific inversion between humans and macaques, we first selected all of the non-human primate species included in the multiple alignments of 99 vertebrate genomes against the human genome (GRCh38/hg38) in the UCSC Genome Browser. Bushbabies were excluded due to an incomplete assembly. Hence, ten non-human primate species were retained for downstream analyses, including chimpanzees (panTro6), gorillas (gorGor6), orangutans (ponAbe3), gibbons (nomLeu3), crab-eating macaques (macFas5), baboons (papAnu4), green monkeys (chlSab2), marmosets (calJac4), squirrel monkeys (saiBol1) and rhesus macaques (rheMac10Plus).

We downloaded the Multiple Alignment Format (MAF) file derived from syntenic net alignment for each non-human primate species from the UCSC Genome Browser. For crab-eating macaques, the MAF file was not available in the UCSC Genome Browser, so we manually generated it using netToAxt. For rhesus macaques, we used the MAF file against hg38, which was constructed from scratch as described in the section “**Detection of inversions between humans and macaques**”. We partitioned the genomic context of each inversion into three parts: the inversion body itself and the upstream and downstream flanking regions, which were twice as long as the inversion. For each inversion, basewise alignments of these three parts were then extracted from the MAF file, and the alignment status of each part was determined if the following criteria were met: 1) 50% of bases fell on the same chromosome, and 2) 80% of them were oriented in the same direction. The regions were defined as inverted if the following criteria were met: 1) at least one flanking region fell on the same chromosome as the inversion body region, and 2) these two regions were aligned in the opposite direction with respect to the human genome. Notably, inversions were removed from downstream ancestor state reconstruction if they were not detected as inverted under these criteria between humans and rhesus macaques, as mentioned above.

Next, we used Phytools (version: 1.0-1)(72) to infer the ancestral states of these inversions, combining the inversion state across humans and other species and the primate phylogeny information. The phylogenetic trees of the primates were extracted under the primate node in the phylogenetic tree used in the MULTIZ multiple alignment against GRCh/hg38. We performed 1,000 stochastic character mappings under the ER model (equal rates for all permitted transitions) for each inversion. For regions with ambiguous inversions, we set equal prior probabilities of 0.5 for both states. Then, we summarized these stochastic maps by determining the posterior probability at each node and uncertain tip, and the state with a posterior probability greater than 0.6 was defined as the estimated state.

With the estimated states at nodes and tips, we then categorized these inversions into two main classes of lineage specificity based on their common ancestors, including Hominoidea-origin and Cercopithecidae-origin sets. Human-specific and macaque-specific inversions were further highlighted from these two sets of distinct origins. For instance, an inversion was classified into a Hominoidea-origin lineage if it was derived in the ape branch and absent from any other nodes or tips. Notably, those inversions that failed in ancestral inference due to uncertainty at some nodes or tips were labeled as ambiguous and were excluded from the subsequent analyses.

### Polymorphic status of human-specific inversions

To assess the polymorphic status of candidate human-specific inversions, we first gathered the genome sequences of 15 public human genome assemblies, including individuals of multiple distinct ethnicities or regions. Most of them were *de novo* assembled using long-read sequencing strategies. Notably, three assemblies (mHomSap3, HG00733, and HuRef) were diploid and were split into maternal and paternal haplotypes for downstream analyses. The metadata of these assemblies can be found in **Table S10**. Similar to the method for the detection of inversions between humans and macaques, we performed pairwise, whole-genome alignment for each assembly against hg38 using DoBlastzChainNet.pl. The output MAF alignment files were used to determine the form of each candidate human-specific inversion in these human assemblies using the same criteria indicated in the previous section. An inversion was defined as fixed if all individuals exhibited concordant alignment with hg38 in this region.

We then evaluated the polymorphic status of the orthologous regions of these candidate human-specific inversions in the population of 572 macaques. Specifically, high-quality (mapping quality > 30), improperly aligned, paired-end reads were extracted around the breakpoints of each candidate inversion, which were normalized according to the length of the regions and the total sequencing depth. Inversions identified with a high allele frequency (frequency > 0.5) in the macaque population were used as the positive control. Two sets of randomly shuffled genomic regions with the same length and chromosome distribution were used as the negative controls. The normalized, improperly aligned, paired-end reads were then calculated to assess the status of each region in each macaque animal. An inversion was defined as polymorphic in the macaque population when at least one animal showed inverted alignment according to the rheMac10Plus genome assembly.

To calculate the genetic divergence accumulated in the human or macaque lineage after the divergence of the species, the ancestral sequences were inferred from the EPO multiple alignments of ten primates (bonobo, chimpanzee, crab-eating macaque, gibbon, gorilla, human, macaque, mouse lemur, Sumatran orangutan, vervet-AGM; **Fig. S11**). Regions downstream and upstream of the inverted regions with matched lengths were used as the controls.

### Annotation and classification of fixed human-specific inversions

The features of a “strong effect inversion” include larger size (> 10 kb), changing gene structures, overlap with regulatory elements (promoters/enhancers), and changing chromatin architectures of compartments, TADs and loops in brain (fetal CP, GZ and adult PFC). An inversion was defined as a strong effect inversion if it showed at least one of these features; otherwise, it was classified as a weak-effect inversion. The inversions were ranked according to their combined effects on gene structure or expression regulation.

### Gene expression analyses

The RNA-seq reads of human and rhesus macaque across organs and developmental stages were downloaded from Cardoso-Moreira *et al.*(73). The FASTQ files were aligned against GRCh38/hg38 genome for human samples and rheMac10Plus genome for rhesus macaque samples using HISAT2 (version: 2.2.1)(74). PCR duplicates were removed using Picard (version: 2.25.4). To minimize the discrepancy of gene annotation across species, we used XSAnno (version: 1.0)(75) to generate the comparative gene annotations between human and rhesus macaque based on GENCODE v33. The read numbers mapped to each gene were counted using featureCounts (version: 2.0.2)(76). DESeq2 (version: 1.26.0)(77) was used to perform differential expression analysis and the log_2_-transformed fold change (human vs. macaque comparison) derived from it for orthologous genes was used to measure the expression divergence. Genes resided in inversions and 50% flanking regions were defined as putatively inversion-associated genes. We used processed RPKM (Reads Per Kilobase per Million mapped reads) tables of human, rhesus macaque and mouse samples for quantification of gene expression levels, which were also obtained from Cardoso-Moreira *et al*.(73).

### Enrichment analyses of contact losses

A permutation test was performed to quantify the degree of contact loss associated with an inversion. Briefly, we first counted the significant interactions in the 1 Mb flanking regions bridging the inward breakpoint (BP2) in rhesus macaques. Then, the conserved portions of flanking regions were retrieved from the human genome using LiftOver, with the minimum match percent set as 0.7. Next, the significant interactions were counted. We used the extent of loss to quantify the degree of significant interaction loss, which was defined as (Count_m_ – Count_h_)/Count_m_, where Count_m_ and Count_h_ are the significant interactions of flanking regions in macaques and humans, respectively. Next, BEDtools (version: 2.26.0)(78) was used to randomly shuffle intervals 10,000 times for BP2, and the extent of loss was calculated for each permutation following the same approach described above. The empirical P value was then calculated to assess the degree of contact loss.

### Dual-luciferase reporter assay

The *APCDD1* promoter region of human and rhesus macaque were amplified and inserted into the pGL3-Basic plasmid, respectively. The mutant plasmids were constructed by introduction of point mutations using TaKaRa MutanBEST Kit. Before transfection, 293T cells were plated to 80% confluence in 12-well plates. Cells were then co-transfected with the *APCDD1*-promoter-constructs and pRL-TK plasmid expressing *Renilla* luciferase as the internal control. After 24h, the luciferase assay was performed using the dual luciferase reporter assay system (Promega) according to the manufacturer’s instructions. *Firefly* and *Renilla* luciferase activities were measured by luminometers.

### Human embryonic stem cell (hESC) culture and neural induction

hESCs (H9 cells) were cultured in Essential 8 Medium (Thermo Fisher Scientific, A1517001) on Matrigel (Corning) in 6-well plates. The cells were regularly tested for mycoplasma. Neural induction was performed using small molecules as previously described(79). In brief, hESCs were detached with dispase to form embryoid bodies (EBs) in neural induction medium (NIM) with 100 nM LDN193189 and 10 μM SB431542. The medium was changed every day. After four days, the EBs were plated in neural progenitor medium (NPM) with 100 nM LDN193189 and 10 μM SB431542 onto plastic plates coated with growth factor-reduced (GFR) Matrigel. After 8 days of induction, the hESCs were differentiated into human NPCs.

The NPCs were maintained using NPM with 10 ng/ml basic fibroblast growth factor (bFGF) on GFR Matrigel. The culture medium was changed every day, and the cells were passaged every six days with Accutase.

### Plasmid construction and lentivirus production

The full-length *APCDD1* sequence was amplified from the cDNA of 293T cells and cloned into the PCDH-CAG-IRES-mCherry backbone (from the laboratory of Dr. Baoyang Hu). Lentiviruses were produced by transfecting 293T cells using the PCDH-*APCDD1*-IRES-mCherry plasmid or PCDH-IRES-mCherry plasmid combined with the packaging plasmids psPAX2 and pMD2.G. Then, the supernatant was collected for three days after transfection and concentrated with a filter device (Amicon Ultra15 Centrifuge Filters, Merck, Cat# UFC910008).

### Lentiviral infection of NPCs

Human NPCs were plated on Matrigel-coated coverslips in 24-well plates with NPM plus 5 μM ROCK inhibitor two days before viral infection. Then, a concentrated lentivirus solution was added to the NPM with 5 μM ROCK inhibitor to infect cells. The lentivirus medium was removed on the second day. After infection, the cells were cultured with NPM without bFGF, and the medium was changed every two days for differentiation. Five days, ten days and 15 days after viral infection, cells grown on coverslips were collected and fixed with 4% PFA for 30 min at room temperature. Then, the PFA was washed with PBS three times. The fixed cells were conserved in PBS at 4 °C for immunofluorescence staining.

### Immunofluorescence staining

The coverslips were incubated for 1 h in blocking buffer (1× PBS, 5% (w/v) BSA, 3% (v/v) Triton X-100) and incubated with primary antibodies diluted in 1% (w/v) BSA and 1% (v/v) Triton X-100 in PBS at 4 °C overnight. Antibodies including anti-mCherry, anti-SOX2, and anti-TUJ1 were used. Then, the coverslips were washed with PBS three times and incubated with Alexa Fluor secondary antibodies diluted in the same solution used for the primary antibodies at room temperature for 1.5 h. After three PBS washes, the coverslips were stained with Hoechst diluted 1/1,000 in PBS for 15 min. Then, the coverslips were washed with PBS and mounted on glass slides with mounting reagent. Imaging was then performed using a Zeiss LSM980 confocal microscope, and the images were processed with ZEN software. For statistical analyses, cells were manually counted using ImageJ(80). At least 20 images were used for counting in every experimental condition.

### Construction of *Apcdd1*-deficient mice

According to the structure of the *Apcdd1* gene, the region from exon 2 to exon 3 of the *Apcdd1*-203 (ENSMUST00000236135.1) transcript was defined as the knockout region. We used CRISPR/Cas9 technology to modify the *Apcdd1* gene. Briefly, sgRNA was transcribed *in vitro*. Cas9 and sgRNA were microinjected into fertilized eggs of C57BL/6J mice, and the fertilized eggs were transplanted to obtain positive F0 mice. A stable F1 generation mouse model was obtained by mating positive F0-generation mice with C57BL/6J mice. *Apcdd1*^+/+^ (WT), *Apcdd1*^+/-^ (Het) and *Apcdd1*^-/-^ (DKO) mice were then generated using three different breeding schemes: Het x Het, Het x WT, and Het x DKO.

### Western blottings

Total protein was extracted from the mouse brain using RIPA lysis buffer (Solarbio R0020) supplemented with 1 mM PMSF and complete protease inhibitor cocktail. Protein concentrations were determined using a Pierce BCA Protein Assay (Thermo Fisher 23227). The primary antibodies were anti-Apcdd1 (dilution: 1:1,000; NOVUS; NB110-92756), anti-Apcdd1 (dilution: 1:1,000; R-D; MAB10501-100), and anti-GAPDH (dilution: 1:1,000; Abcam; ab8245). The secondary antibodies were HRP-conjugated goat anti-rabbit IgG (EARTH, E030120-01) and HRP-conjugated goat anti-mouse IgG (EARTH, E030110-01).

### scRNA-seq in mouse embryonic brains

The E10.5 mouse embryos were dissected from pregnant mice sacrificed by anesthesia and decapitation. The head part of embryo was dissected out by fine tweezers and stored in Shbio®Tissue Preservation Solution (21903-10) overnight, which were then dissociated using Shbio®Tissue Dissociation Kit (219517-10). Briefly, tissues were incubated in 1 ml enzyme mix on a metal rotor at 37 °C for 15 min, during which specimens were gently mixed for 20 times using 1,000-ml tips every 7 min. Cell dissociations were terminated by adding 2 ml of complete medium and then suspensions were filtered through a 100-µm strainer and centrifuged for 10 min at 500g. The supernatant was discarded and the cell pellet was resuspended with 300 µl suspension buffer (1X DPBS containing 2% FBS). Dyed by 50 ng/ml DAPI for 3 min and refiltered with a 40-µm strainer, cell suspensions were then sorted with FACS BD Aria III sorter to remove dead cells, fragments and lumps. Cell numbers of suspensions were measured with fluorescence cell counter (Luna-FLTM, Logos biosystems) and hemocytometer using trypan blue staining. Cell suspensions were then adjusted to a proper cell concentration for library construction of scRNA-seq.

scRNA-seq was then performed using the DNBelab C high throughput single cell RNA library prep kit (MGI) following the manufacturer’s protocol. The QubitTM Flex Fluorometer (Thermo Fisher Scientific) and Agilent Fragment Analyzer were used to determine the concentration and assess the quality of the libraries, which were then subjected to DNBSEQ-2000RS for sequencing.

Cell barcode and unique molecular identifiers (UMI) sequences were parsed using DNBelab_C_Series_HT_scRNA-analysis-software. Raw reads were then aligned against GRCm38/mm10 genome and GENCODE M25 gene annotation using *DNBC4tools run* with default parameters to generate sparse gene count matrices. Scanpy (version: 1.9.1)(81) was used for downstream analyses. Low-quality cells were filtered out considering the abnormal gene count and high mitochondrial read percentage. We normalized the gene expression by scaling total UMIs to 10,000 in each cell followed by log transformation. The BBKNN algorithm(82) was used to remove batch effects across four samples. Then, principal component analysis (PCA) was performed and the top 50 principal components (PCs) were used for uniform manifold approximation and projection (UMAP) and Leiden clustering(83). We then determined differentially expressed genes using Wilcoxon rank-sum test method for each cluster. The known marker genes(32–36) was used to assign cell type identities.

## Data and code availability

The PacBio sequence data, Bionano optical map, Illumina whole-genome sequencing reads, Hi-C sequencing data and scRNA-seq have been deposited in NCBI under the project accession number PRJNA748883 and GEO website under accession number GSE213693 and are publicly available. Genome assembly rheMac10Plus, SNV and SV datasets in the rhesus macaque population have been posted to the website of RhesusBase (URL: https://rhesusbase.com/download.jsp). The code employed in this study has been deposited in GitHub under https://github.com/foggieding/1000MonkeyGenome.

## Author Contributions

C.Y.L. and L.Z. conceived and designed the study. W.D., X.L., and J.Z. analyzed most of the data and performed statistical analysis. X.Z., J.W., C.X., T.L., Q.Y., J.C.Y., J.Q., L.T., and X.X. analyzed part of the data. M.J. performed the NPC overexpression experiments. M.J., C.L., X.L., and Q.Y. performed the mouse experiments. X.L. and M.J. performed the dual-luciferase reporter assays. M.Z. performed the FISH experiments. Y.C., F.M., B.Z., J.Q., Q.P., Q.Z.Z., Z.L., A.F., X.Z., J.J.Z., Y.S., B.H. and N.A.A. contributed to data interpretation. C.Y.L., L.Z., W.D., X.L., and J.Z. wrote the paper. All authors read and approved the final manuscript.

## Competing Interest Statement

The authors declare no competing interests.

## Acknowledgments

We thank Dr. Yong E. Zhang at the Institute of Zoology, Chinese Academy of Sciences for insightful suggestions. We thank Ms. Shuang Gao at MGI Tech for technical support. We thank the National Center for Protein Sciences at Peking University in Beijing, China, for assistance with the DNA FISH experiment. This work was supported by grants from the Ministry of Science and Technology of China (National Key Research and Development Program of China, 2018YFA0801405 and 2019YFA0801801), the National Natural Science Foundation of China (31871272), the Chinese Institute for Brain Research (2020-NKX-XM-11), and the Clinical Medicine Plus X – Young Scholars Project, Peking University (PKU2022LCXQ015).

## Notes

### Competing Interest Statement

The authors have declared no competing interest.

