## Supplementary Information for "Adaptive Functions of Structural Variants in Human Brain Development"

### Supporting Information for
### Adaptive Functions of Structural Variants in Human Brain Development

Wanqiu Ding[1,10], Xiangshang Li[1,10], Jie Zhang[1,10], Mingjun Ji[1,10], Mengling Zhang[3], Xiaoming Zhong[1], Yong Cao[4], Xiaoge Liu[1], Chunqiong Li[2], Chunfu Xiao[1], Jiaxin Wang[1], Ting Li[1], Qing Yu[1], Fan Mo[5], Boya Zhang[5], Jianhuan Qi[5], Jie-Chun Yang[1], Juntian Qi[1], Lu Tian[1], Xinwei Xu[1], Qi Peng[1], Wei-Zhen Zhou[6], Zhijin Liu[7], Aisi Fu[8], Xiuqin Zhang[1], Jian-Jun Zhang[9], Yujie Sun[3], Baoyang Hu[5], Ni A. An[1], Li Zhang[2,*] and Chuan-Yun Li[1,2,*]

[1]Institute of Molecular Medicine, College of Future Technology, Peking University, Beijing, China
[2]Chinese Institute for Brain Research, Beijing, China
[3]State Key Laboratory of Membrane Biology, Biomedical Pioneer Innovation Center (BIOPIC), School of Life Sciences, Peking University, Beijing, China
[4]Department of Neurosurgery, Beijing Tiantan Hospital, Capital Medical University, 119S Fourth Ring Rd W, Fengtai District, Beijing, China
[5]State Key Laboratory of Stem Cell and Reproductive Biology, Institute of Stem Cell and Regeneration, Institute of Zoology, Chinese Academy of Sciences, Beijing, China
[6]State Key Laboratory of Cardiovascular Disease, Fuwai Hospital, Chinese Academy of Medical Sciences and Peking Union Medical College, Beijing, China
[7]College of Life Sciences, Capital Normal University, Beijing, China
[8]Wuhan Dgensee Clinical Laboratory, Wuhan, China
[9]CAS Key Laboratory of Mental Health, Institute of Psychology, Beijing, China
[10]These authors contributed equally: Wanqiu Ding, Xiangshang Li, Jie Zhang, Mingjun Ji

*Corresponding authors: Chuan-Yun Li and Li Zhang
 (C.Y.L.), (L.Z.)

**This PDF file includes:**

Figures S1 to S15
Tables S1 to S3, S6, S8, S10 to S11
Legends for Tables S4 to S5, S7, S9, S12

**Other supporting materials for this manuscript include the following:**

Tables S4 to S5, S7, S9, S12

#### Supplementary Figures

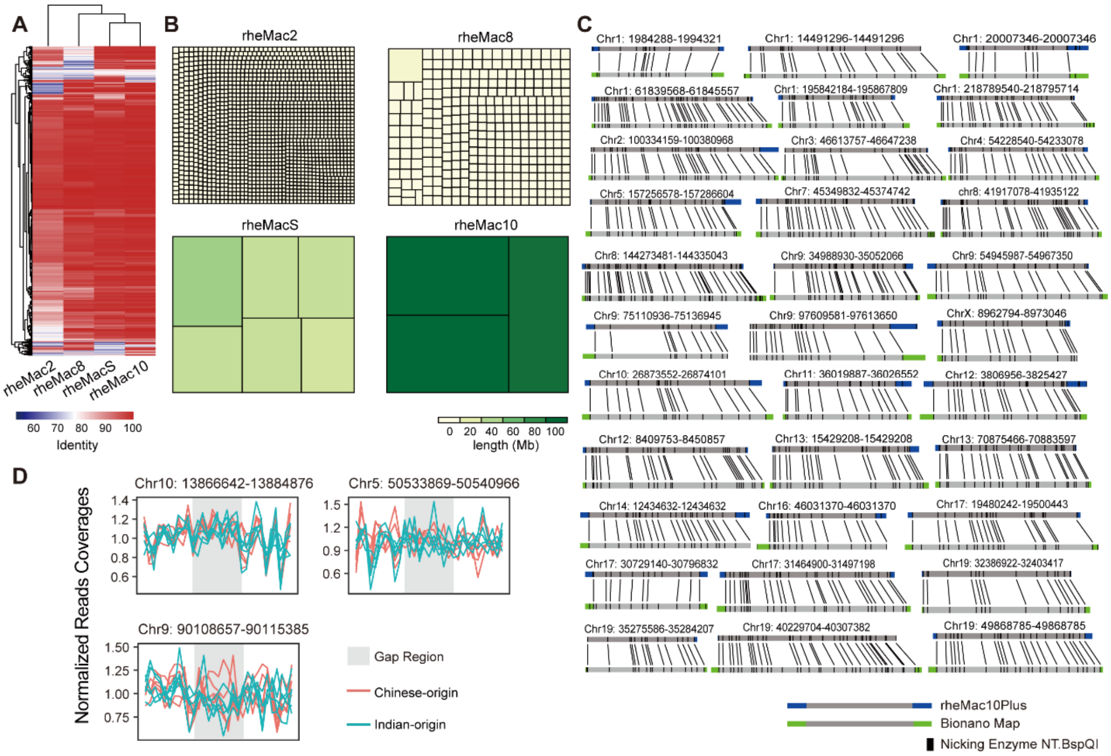


**Fig. S1. Gap closure of the macaque reference genome.** (**A**) Heatmap showing the base identity of four independent assemblies of the macaque genome (rheMac2, rheMac8, rheMacS and rheMac10), as evaluated by BAC sequences. (**B**) Treemaps showing the different distributions of contig sizes in the four assemblies of the macaque genome. The rectangles in each genome assembly represent the largest contigs accounting for 200 Mb of the assembly. (**C**) Visualization of the alignments between Bionano optical maps and the improved macaque reference genome (rheMac10Plus). For the gap regions and their flanking regions, the enzyme cleavage sites identified from the optical map or defined based on the reference genome sequences are shown with black bars, and the continuous and correctly spaced alignments between them are connected by black lines. (**D**) Reads coverages of the filled gaps and their flanking regions, according to the genome sequencing of ten macaques (orange: Chinese-origin macaques; green: Indian-origin macaques. Gap regions are indicated by gray shaded rectangles).

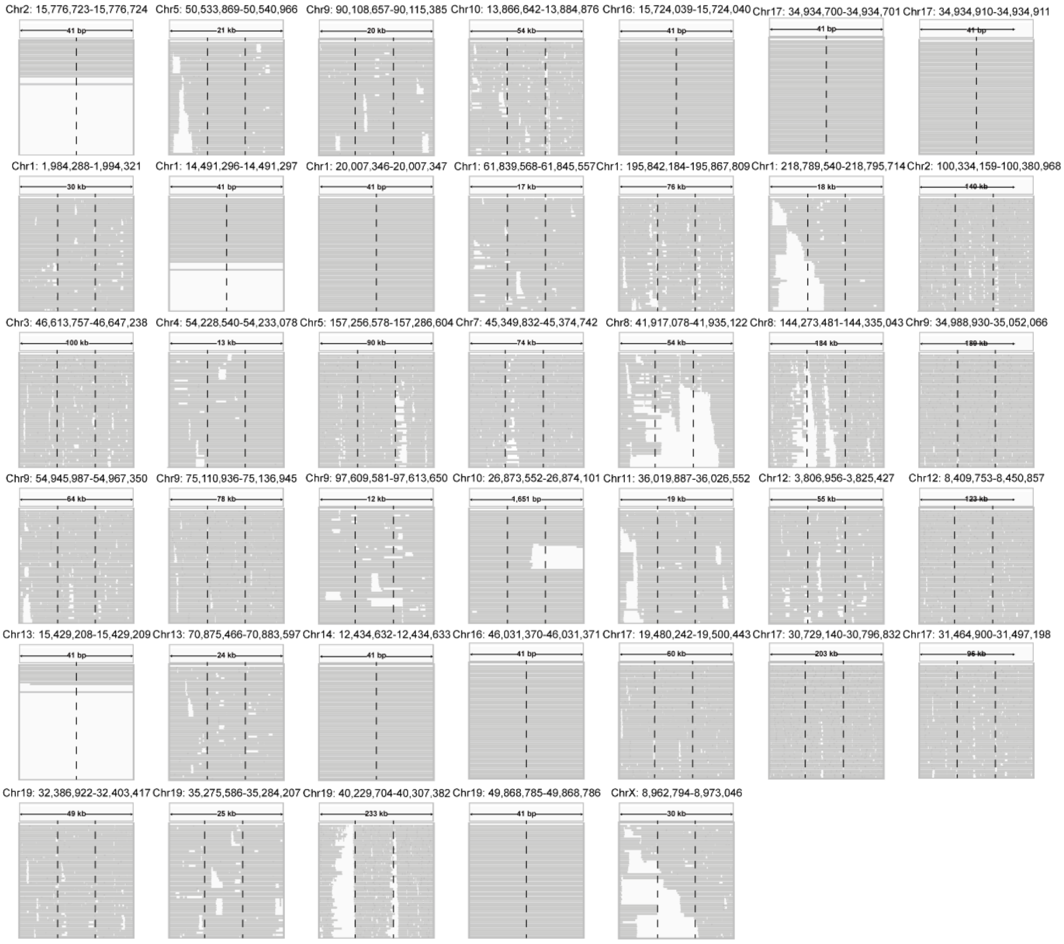


**Fig. S2. Verification of 40 gap closures by long HiFi reads of PacBio sequencing.** For each gap region (indicated by the black dashed lines) and its length-matched, upstream/downstream flanking regions, the long reads covering these regions were aligned and shown.

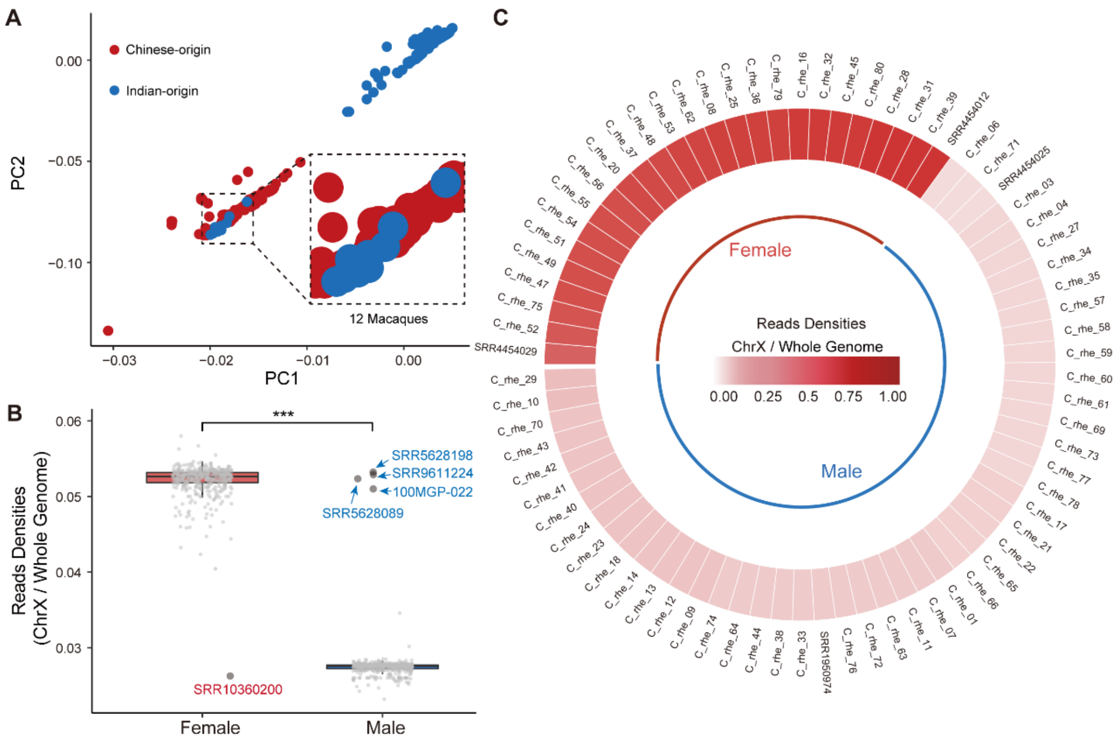


**Fig. S3. Revision of the source and sex information of macaques.** (**A**) PCA plot showing the genetic distances and relationships of the 1,026 macaques, which were clustered into two major groups (red: Chinese-origin macaques; blue: Indian-origin macaques). The 12 macaques with confusing resource information in the original report are highlighted in a zoomed-in view. (**B**) The distribution of reads densities on the X chromosomes of 1,026 macaques, normalized with that in the whole genome for the corresponding macaque individual. The IDs of macaques with confusing sex information are indicated. The Wilcoxon rank-sum test was performed. ***P value < 0.001. (**C**) Heatmap showing the reads densities on the X chromosomes of 74 wild-caught Chinese-origin macaques.

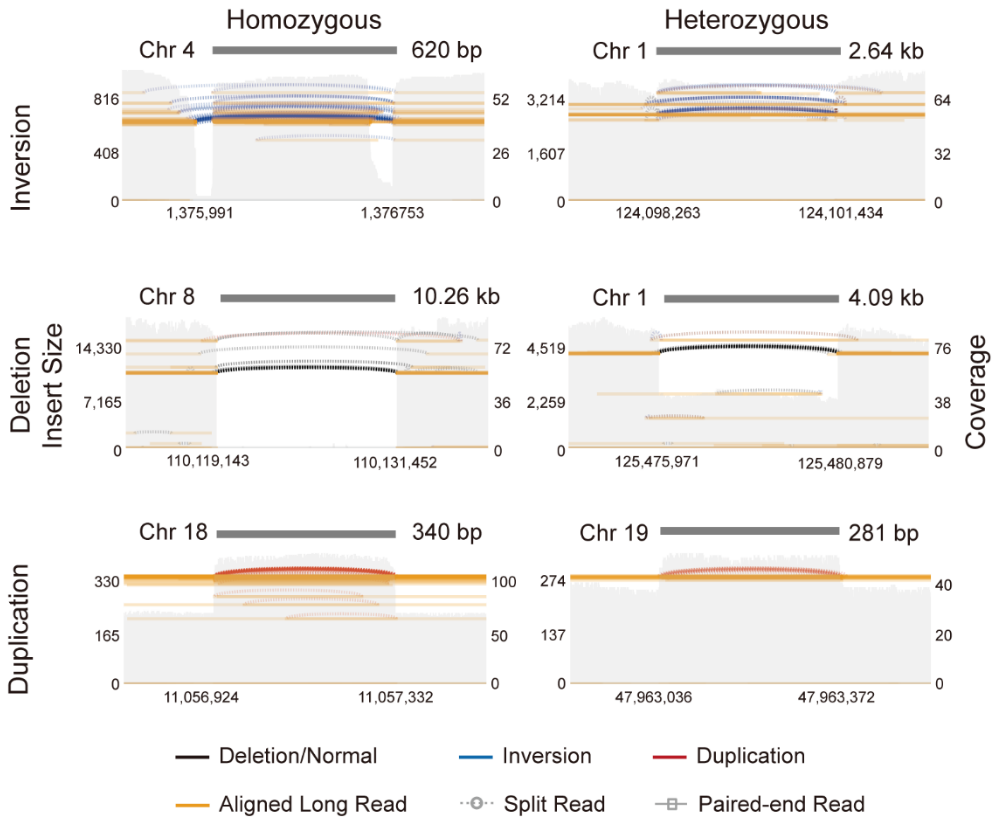


**Fig. S4. Samplots of representative SVs verified by long-reads sequencing.** The aligned long-reads are color-coded according to the type of alignments (concordant/discordant insert size, pair order, split alignment, or long read). The coverage of each region is shown with a gray-filled background.

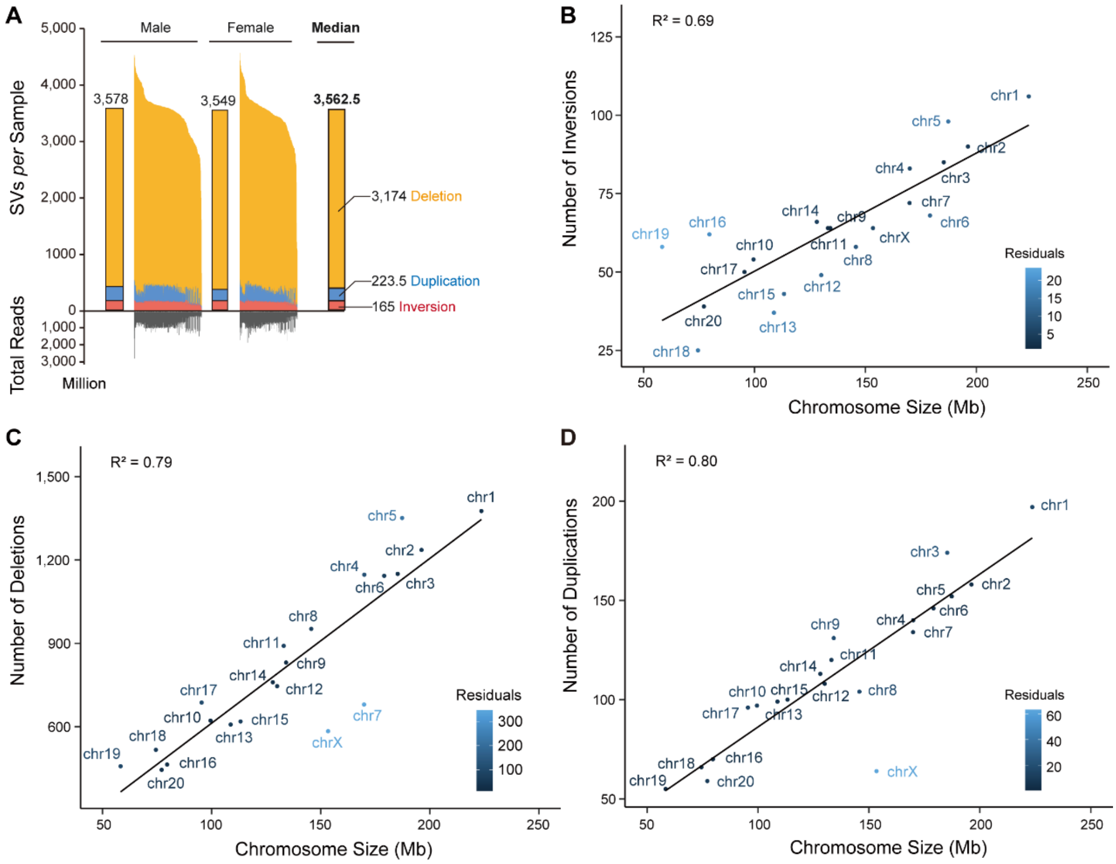


**Fig. S5. Characteristics of the SV map in the macaque population.** (**A**) The distribution of the count of SVs *per* macaque genome for three types of SVs in macaques of different sexes. The median number of SVs is shown for each group. The total number of deep sequencing reads for each macaque is also shown. (**B-D**) Scatter plots showing the correlation between chromosome size (in Mb) and the numbers of inversions (**B**), deletions (**C**) and duplications (**D**) located on each chromosome.

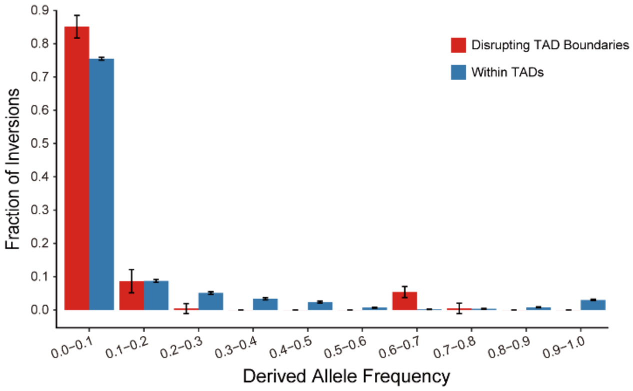


**Fig. S6. Site frequency spectra of inversions with different locations.** Site frequency spectra of the derived alleles for inversions disrupting TAD boundaries (red) and inversions within TADs (blue), indicated by Hi-C data of fetal CP.

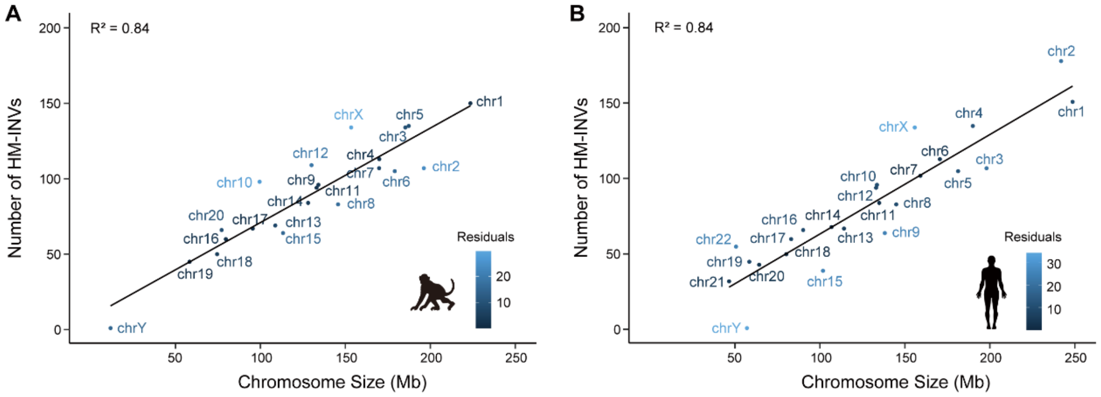


**Fig. S7. Correlations between the length of the chromosome and the number of species-specific inversions between humans and macaques.** (**A-B**) Scatter plots showing the correlation between chromosome size (in Mb) and the number of HM-INVs located on each chromosome in rhesus macaque (**A**) and human (**B**).

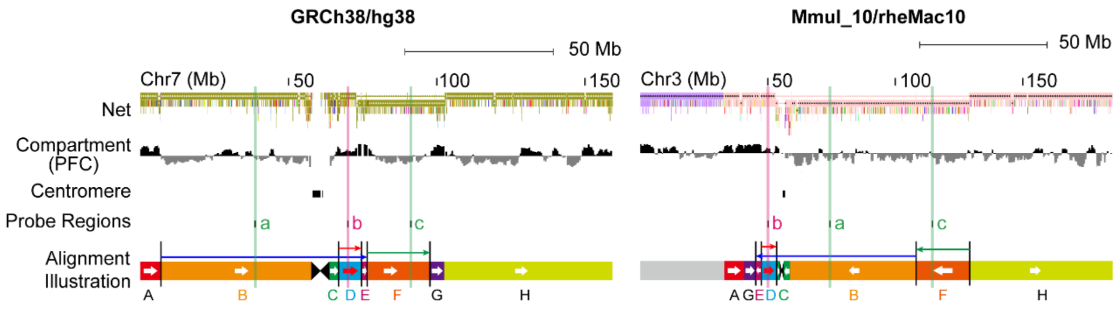


**Fig. S8. Probe design for FISH validation.** Schematic diagram showing the probe design for FISH assays performed to clarify the structures of one complex HM-INV in humans (left) and macaques (right). The Net track shows the alignment between the human and macaque genomes, followed by the information for A/B compartments and the positions of the centromeres. The locations of the three probes designed (a, b and c) are shown in the two panels, with the same colors used in **Fig. 4D**. The pairwise alignment between the two genome assemblies is illustrated in color-coded blocks from A to H, with the inversions indicated by the arrows.

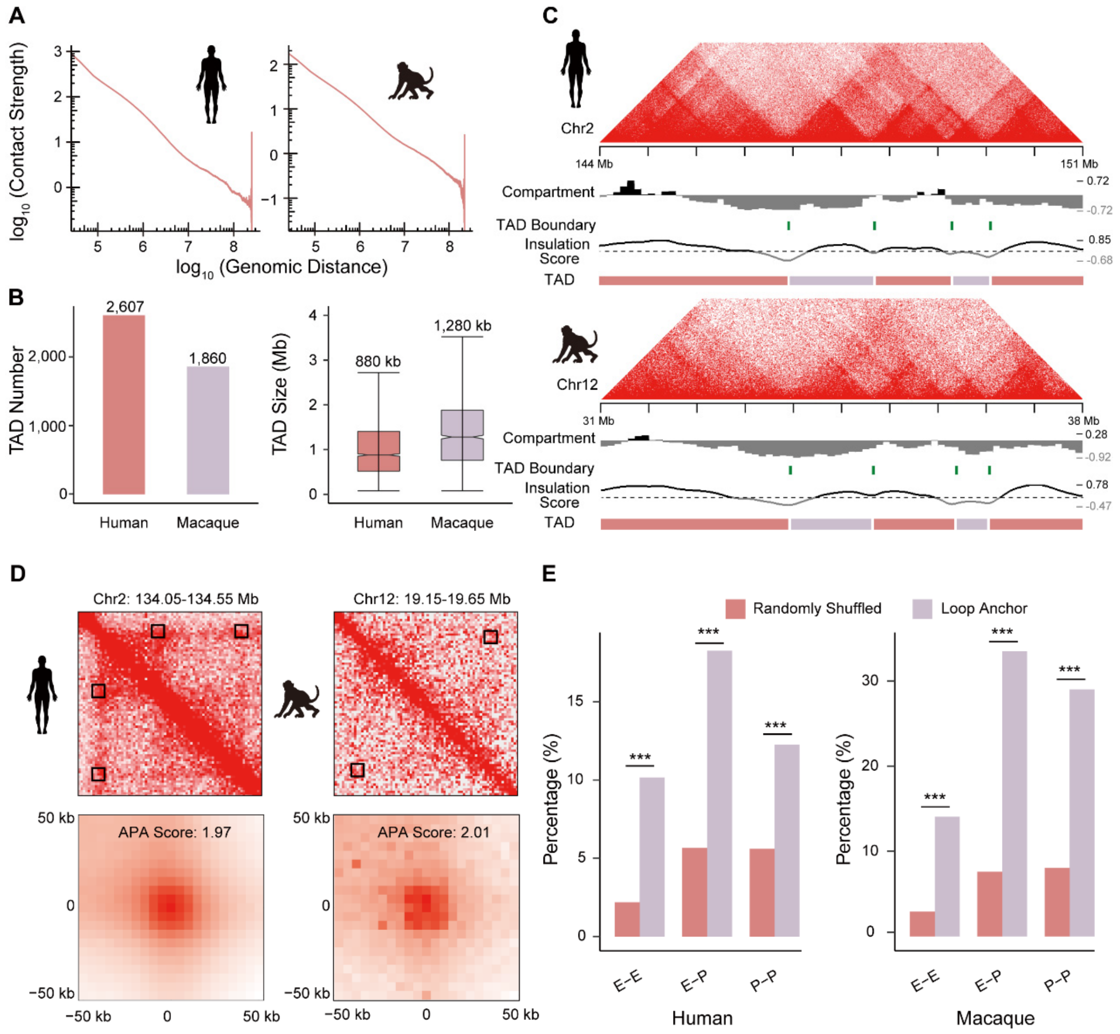


**Fig. S9. High-resolution Hi-C contact maps in human and macaque.** (**A**) Contact strength as a function of the $\log_{10}$-transformed genomic distance in human and macaque genomes. (**B**) The numbers (left) and sizes (right) of the TADs identified in human and macaque. (**C**) Representative Hi-C contact maps in human (top) and macaque (bottom). For each region, the information of the A/B compartments is shown. The TADs are shown in red and purple blocks, and the positions of the TAD boundaries are highlighted with green bars. Genomic regions (in 40 kb windows) with positive insulation scores are represented by black lines, whereas the regions with negative scores are represented by gray lines. (**D**) Representative loop views (top) for one position in human (left) and another position in rhesus macaque (right), with loops shown in dashed boxes. Aggregate peak analysis plots (APA, bottom) showing the combined signals for all of the loops, as detected in human (left) and rhesus macaque (right). (**E**) The percentage of enhancer-enhancer (E-E), enhancer-promoter (E-P) and promoter-promoter (P-P) interactions for loop anchors and randomly shuffled regions in human and rhesus macaque. Fisher's exact test was performed. ***P value < 0.001.

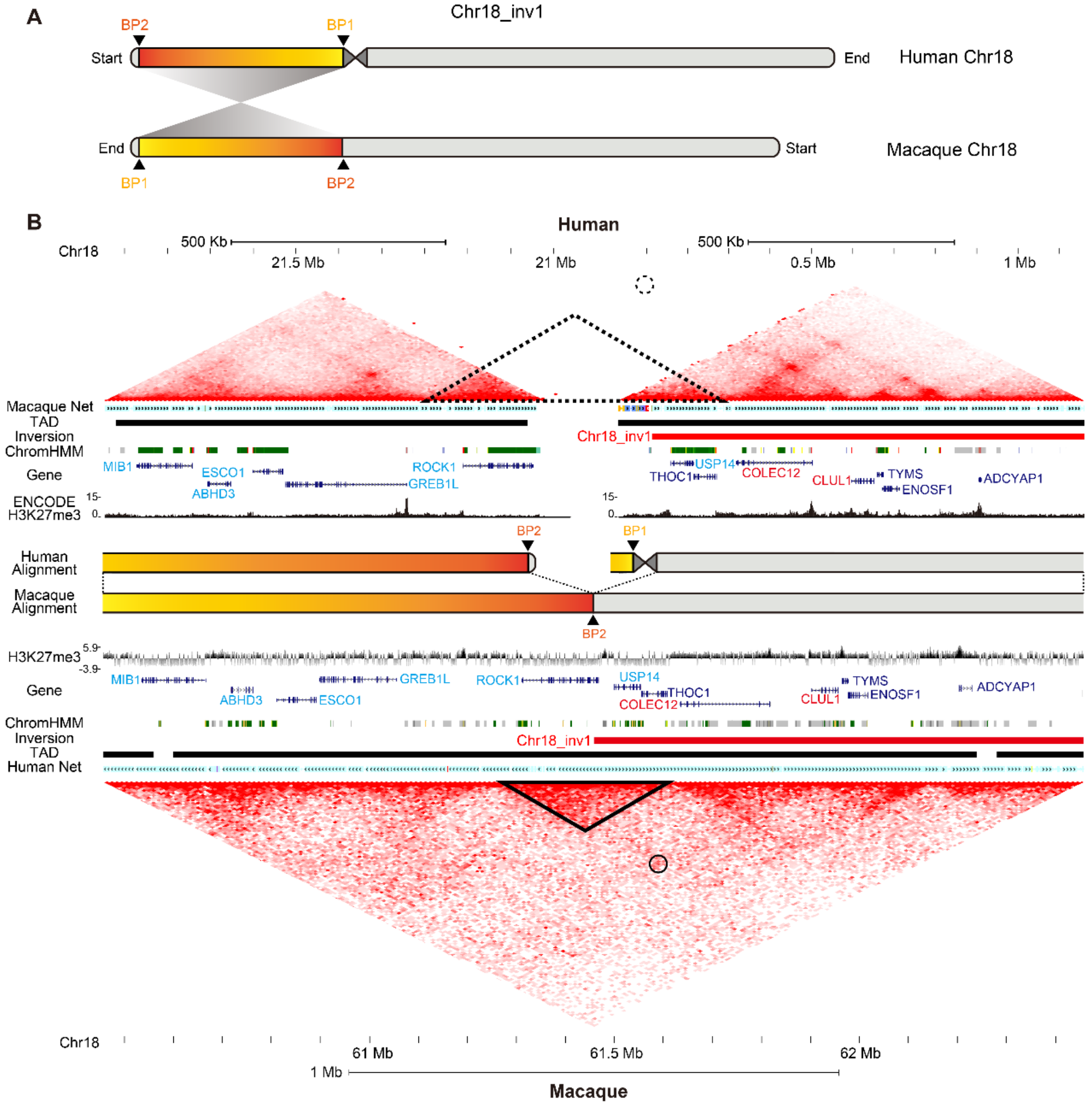


**Fig. S10. Hi-C contact maps in human and macaque for a human-specific inversion.** (**A**) Schematic illustration of a human-specific inversion on chromosome 18. (**B**) The contact map for the human-specific inversion in human (top) and the orthologous region in macaque (bottom). The interactions predicted to be lost as a consequence of the inversion are indicated by black triangles and circles. The human-macaque net tracks are shown to indicate the inverted alignment. The TADs are represented by black blocks. The chromatin states are indicated in the ChromHMM and H3K27me3 tracks. In comparison with the expression in macaque brains, the genes downregulated in humans were shown in blue, while the upregulated genes were shown in red. A schematic diagram of the detailed alignment between human and macaque genomes is also shown.

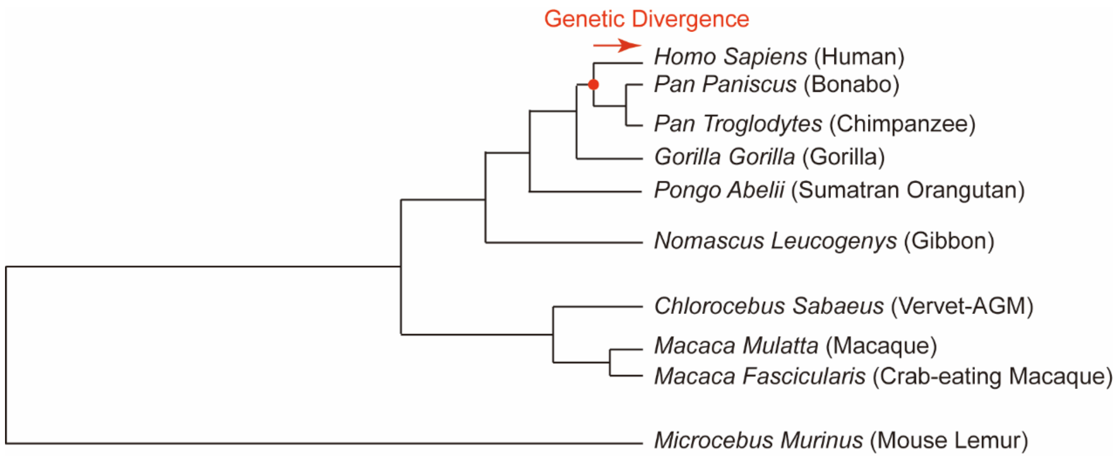


**Fig. S11. Tracing the ancestral state with the EPO pipeline.** Schematic diagram of the evolutionary tree of ten primate species in the EPO pipeline to trace the ancestral state of variants. The node of the common ancestor of humans and chimpanzees is highlighted with a red dot, and the branch used to calculate genetic divergence in **Fig. 5** is indicated with a red arrow.

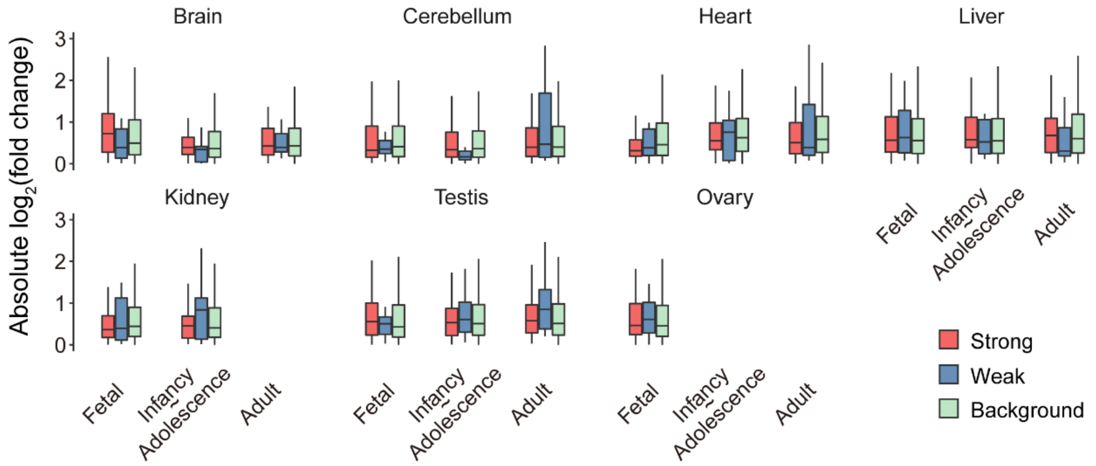


**Fig. S12. Comparisons of gene expression in human and macaque tissues.** $Log_2$-transformed fold changes in gene expression between human and macaque tissues across various developmental stages for strong effect inversions (Strong), weak effect inversions (Weak) and genome-wide ortholog pairs as a background (Background).

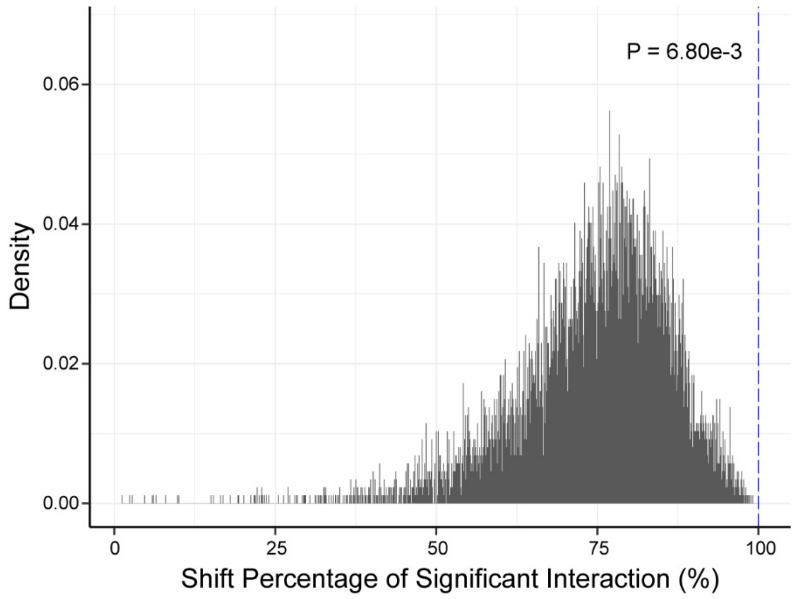


**Fig. S13. Contact loss at chr18_inv1.** Histogram showing the distribution of the "shift of significant interactions" in 10,000 permutations of randomly shuffled breakpoints, with the blue dotted line indicating the value of the breakpoint downstream of this inversion (chr18_inv1, BP2). A permutation test was performed to calculate the P value.

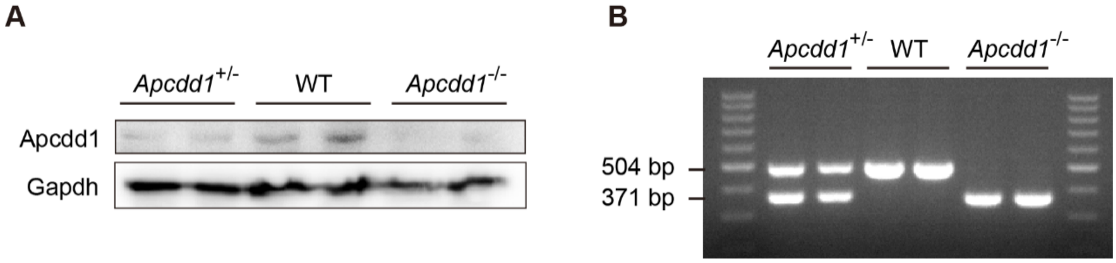


**Fig. S14. Evaluation of knockout efficiencies in *Apcdd1*-deficient mice.** (**A**) Western blots showing the protein expression of Apcdd1 and Gapdh in the brains of wild-type (WT) and *Apcdd1*-deficient mice (*Apcdd1*$^{+/-}$ and *Apcdd1*$^{-/-}$). (**B**) PCR-amplified products from wild-type (504 bp) and *Apcdd1*-deficient mice (*Apcdd1*$^{+/-}$ and *Apcdd1*$^{-/-}$). The bands of 504 bp and 371 bp indicate the PCR products encoded by wild-type *Apcdd1* and mutant *Apcdd1*, respectively.

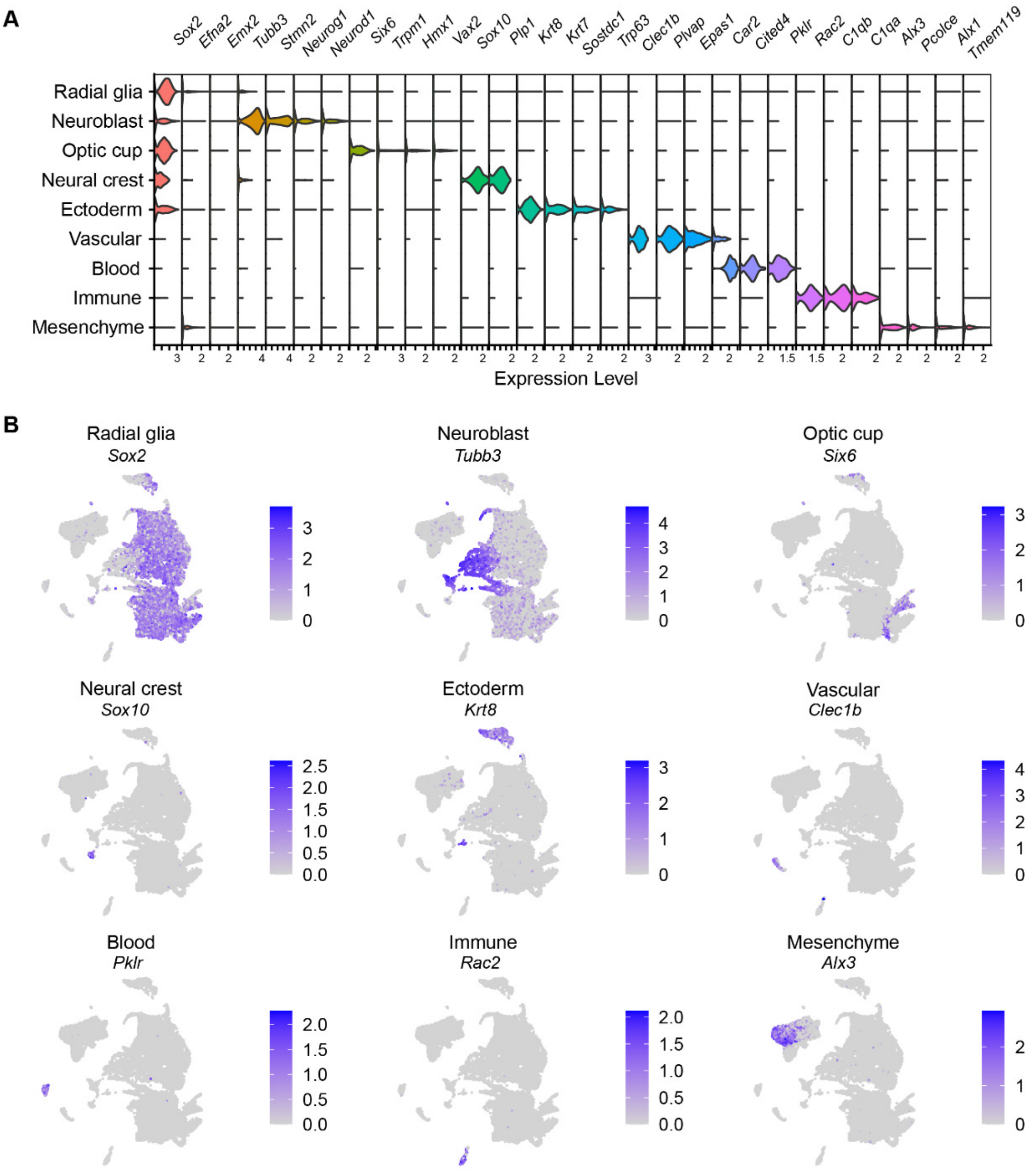


**Fig. S15. Representative marker genes of scRNA-seq from E10.5 mice brains.** (**A**) Violin plot of gene expression levels of select marker genes (columns) for each cell cluster (row). (**B**) Representative markers of scRNA-seq clustering results for each cell type.

#### Supplementary Tables

**Table S1. Statistics of the Bionano optical map data for one captive Chinese-origin macaque.**

| Length Bin | Molecule Count | Quantity (Gb) | Average Length (kb) | Molecule N50 (kb) | Labels (/100 kb) |
|---|---|---|---|---|---|
| **>150 kb** | 1,606,066 | 477.2 | 297.11 | 317.43 | 9 |

**Table S2. Statistics of the whole-genome sequencing data of five Indian- and five Chinese-origin macaques used in the evaluation of gap closure.**

| Macaque ID | Origin | Sex | Sources | Total Reads | Mapped Reads | Total Mapping Rate (%) | Depth |
|---|---|---|---|---|---|---|---|
| **SRR1944102** | Captive Chinese-origin | Female | Xue *et al*. | 1,368,158,572 | 1,364,841,341 | 99.76 | 46.5058 |
| **SRR1944106** | Captive Chinese-origin | Female | Xue *et al*. | 971,523,728 | 965,468,360 | 99.38 | 33.0236 |
| **SRR1952123** | Captive Chinese-origin | Female | Xue *et al*. | 1,245,697,984 | 1,235,247,094 | 99.16 | 42.3432 |
| **SRR1950974** | Wild-caught Chinese-origin | Male | Xue *et al*. | 817,422,924 | 814,214,420 | 99.61 | 34.3879 |
| **SRR1951000** | Wild-caught Chinese-origin | Female | Xue *et al*. | 693,209,140 | 691,446,716 | 99.75 | 29.1624 |
| **SRR7865677** | Captive Indian-origin | Female | Bimber *et al*. | 688,066,936 | 686958223 | 99.84 | 34.7353 |
| **SRR7865679** | Captive Indian-origin | Female | Bimber *et al*. | 746,018,234 | 744,766,728 | 99.83 | 37.6608 |
| **SRR7865683** | Captive Indian-origin | Male | Bimber *et al*. | 751451034 | 750,233,111 | 99.84 | 37.9351 |
| **SRR7865698** | Captive Indian-origin | Male | Bimber *et al*. | 556,988,368 | 555,880,630 | 99.8 | 28.1181 |
| **SRR8474135** | Captive Indian-origin | Male | Bimber *et al*. | 1,095,849,906 | 1,087,253,888 | 99.22 | 37.2496 |

**Table S3. Statistics of PacBio HiFi reads for eight rhesus macaque samples.**

| Sample ID | HiFi Reads | HiFi Yield (bp) | HiFi Read Quality (median) | HiFi Read N50 |
|---|---|---|---|---|
| **920201** | 496,251 | 9,047,694,354 | 31 | 18,532 |
| **031024** | 423,194 | 8,305,659,930 | 31 | 20,601 |
| **97236** | 359,142 | 6,765,613,234 | 31 | 19,266 |
| | 63,001 | 1,303,569,644 | 30 | 20,544 |
| **97009_3_3** | 116,056 | 1,825,651,416 | 32 | 16,095 |
| **9109_35_1** | 61,779 | 1,044,755,305 | 31 | 17,093 |
| **100MGP-018** | 72,040 | 1,215,420,459 | 31 | 17,505 |
| **090089_3** | 35,080 | 635,061,116 | 31 | 18,604 |
| **100MGP-003** | 27,375 | 495,108,397 | 31 | 18,197 |

**Table S6. Statistics for PacBio genome sequencing of one captive Chinese-origin macaque.**

| Type | Total Bases | Total Bases (Subreads) |
|---|---|---|
| **RSII** | 110,826,954,922 | 101,952,187,859 |
| **Sequel** | 195,530,217,571 | 188,638,334,863 |
| **Total** | 306,357,172,493 | 290,590,522,722 |

| Type | Min Read Length | Max Read Length | Mean Read Length | Read N50 Length | Read Count |
|---|---|---|---|---|---|
| **Total** | 50 | 193,083 | 8,700 | 14,339 | 33,399,602 |

**Table S8. Genomic coordinates of the probes and flanking primers used in the FISH assays.**

| Species | Genomic Region Index | Genomic Coordinates | Flanking Primers |
|---|---|---|---|
| **Human (GRCh38)** | a | chr7:38,505,934-38,966,169 | CATCGAAGCGTGTGGCTACC |
| | b | chr7:69,818,995-70,273,729 | CATCGAAGCGTGTGGCTACC |
| | c | chr7:91,071,663-91,570,490 | TCGTTCCGCATTGACCAATC |
| **Rhesus macaque (Mmul_10)** | a | chr3:74,238,416-74,693,339 | CCTGTGCGGAAATCGCGAGA |
| | b | chr3:49,980,073-50,434,877 | TCGTTCCGCATTGACCAATC |
| | c | chr3:114,399,067-114,898,598 | GGATTGCCGCATGGTTTCCG |

**Table S10. Metadata of 15 human genome assemblies used in this study.**

| Genome Assembly | Sequencing Strategy | Population | Ethnic Group | Sex | Contig N50 | GenBank ID |
|---|---|---|---|---|---|---|
| **Ash1.7** | Illumina HiSeq; Oxford Nanopore MinION; PacBio Sequel | European | Caucasian | male | 34,306,965 | GCA_011064465.1 |
| **ASM1490585v1** | PacBio RSII; Bionano; Illumina HiSeq | East Asian | Japan | male | 23,601,496 | GCA_014905855.1 |
| **HG00514_prelim_3.0** | PacBio RSII | East Asian | Han Chinese (Southern) | female | 29,396,877 | GCA_002180035.3 |
| **HG00733_Phased_Diploid** | PacBio Sequel; Illumina HiSeq HiC | American | Puerto Rican | female | 26,292,878 | GCA_003634875.1 |
| **hg01243.v3.0** | Illumina NovaSeq; Oxford Nanopore; PacBio Sequel; PacBio HiFi | American | Puerto Rican | male | 149,697,505 | GCA_018873775.2 |
| **HS1011_v1.1** | Illumina; PacBio (RS and RSII) | American | USA | male | 373,039 | GCA_001292825.2 |
| **KOREF_S1v2.1** | Oxford Nanopore PromethION; PacBio Sequel II HiFi; Illumina HiSeq | East Asian | Korean | male | 20,046,962 | GCA_020497085.1 |
| **KOREF1.0** | Illumina HiSeq; PacBio | East Asian | Korean | male | 47,865 | GCA_001712695.1 |
| **mHomSap3.mat** | PacBio Sequel I CLR; Illumina NovaSeq; Arima Genomics Hi-C; Bionano Genomics DLS | mixed | mixed | male | 12,076,592 | GCA_016695395.2 |
| **mHomSap3.pat** | PacBio Sequel I CLR; Illumina NovaSeq; Arima Genomics Hi-C; Bionano Genomics DLS | mixed | mixed | male | 13,671,004 | GCA_016700455.2 |

| | | | | | | |
|---|---|---|---|---|---|---|
| **NA12878_prelim_3.0** | PacBio RSII | American | Utah/Mormon | female | 16,821,028 | GCA_002077035.3 |
| **NA19240_prelim_3.0** | PacBio | African | Yoruban | female | 29,140,631 | GCA_001524155.4 |
| **PGP1v1** | Oxford Nanopore PromethION; PacBio Sequel II HiFi; Illumina NovaSeq | European | Caucasian | male | 24,698,710 | GCA_020497115.1 |
| **HuRef** | Sanger dideoxy technology | European | Caucasian | male | 108,431 | GCA_000002125.2 |
| **HuRefPrime** | Sanger dideoxy technology | European | Caucasian | male | 100,385 | GCA_000212995.1 |
| **T2T-CHM13v2.0** | PacBio; Nanopore; Illumina; BioNano; Strand-seq | mixed | major European | mixed | 154,260,000 | GCA_009914755.4 |
| **CSA** | Sanger dideoxy technology | mixed | mixed | mixed | 97,525 | GCA_000365445.1 |

**Table S11. Statistics of genome-wide contact maps for adult human and macaque.**

| Species | Total Reads Pairs | Uniquely Mapped Reads Pairs | Valid Pairs | Valid Pairs with Duplication Removed | Cis-interaction | Trans-interaction | Resolution (bp) |
|---|---|---|---|---|---|---|---|
| **Human** | 1,976,103,225 | 1,373,607,770 | 1,358,484,946 | 961,419,088 | 667,101,577 | 294,317,511 | 2,500 |
| **Macaque** | 1,458,885,556 | 772,771,900 | 756,408,416 | 342,056,553 | 183,623,764 | 158,432,789 | 5,450 |

**Table S4 (separate file). Information for 27 captive Chinese-origin macaques in whole genome sequencing.**

**Table S5 (separate file). Statistics for 1,026 rhesus macaques with whole genome sequencing data.**

**Table S7 (separate file). 1,972 inversions between human and rhesus macaque.**

**Table S9 (separate file). 101 candidate human-specific inversions in human and macaque populations.**

**Table S12 (separate file). Classification of 75 fixed human-specific inversions based on putative effects.**
